# Multi-modal meta-analysis of cancer cell line omics profiles identifies ECHDC1 as a novel breast tumor suppressor

**DOI:** 10.1101/2020.01.31.929372

**Authors:** Alok Jaiswal, Prson Gautam, Elina A Pietilä, Sanna Timonen, Nora Nordström, Nina Sipari, Ziaurrehman Tanoli, Kaisa Lehti, Krister Wennerberg, Tero Aittokallio

## Abstract

A caveat of cancer cell line models is that their molecular and functional profiling is subject to laboratory-specific experimental practices and data analysis protocols. The current challenge is how to make an integrated use of omics profiles of cancer cell lines for reliable discoveries. Here, we carried out a systematic analysis of nine types of data modalities using meta-analysis of 53 omics profiling studies across 12 research laboratories for 2018 cell lines. To account for relatively low consistency observed for certain data modalities, we developed a robust data integration approach that identifies reproducible signals shared among multiple data modalities and studies. We demonstrated the power of the integrative analyses by identifying a novel driver gene, ECHDC1, with tumor suppressive role validated both in breast cancer cells and patient tumors. Extension of the approach identified synthetic lethal partners of cancer drivers, including a co-dependency of PTEN deficient cells on RNA helicases.

**Highlights:**

- A comprehensive meta-analysis of 53 multi-modal omics profiles of >2000 cancer cell lines from 12 research laboratories
- An unexpected lack of consistency between TMT-labelled and non-labelled global proteomic profiles
- A non-parametric approach to integrate omics profiles from multiple laboratories and to identify robust molecular patterns in individual cell lines
- The multi-modal data integration reveals novel drivers and potential therapeutic targets, including ECHDC1 in breast cancers and DDX27 in PTEN mutant cancers.

## Introduction

Cancer cell lines have immensely served the purpose of expanding our understanding of cancer biology, and also accelerated the process of developing new targeted therapeutics(Ben-David et al., 2018; Gillet et al., 2013). Analogous to the patient tumor profiling efforts(Hutter and Zenklusen, 2018; Zehir et al., 2017), high throughput ‘omics’ techniques have enabled a deep molecular and genetic characterization of large panels of human cancer cell lines^5–33^. As a result, a high-resolution molecular portrait of the genome(Barretina et al., 2012; Daemen et al., 2013; Ghandi et al., 2019; Iorio et al., 2016; Klijn et al., 2015; Marcotte et al., 2016; Shankavaram et al., 2009), transcriptome(Barretina et al., 2012; Daemen et al., 2013; Ghandi et al., 2019; Iorio et al., 2016; Klijn et al., 2015; Marcotte et al., 2016; Shankavaram et al., 2009), proteome(Coscia et al., 2016; Gholami et al., 2013; Lapek et al., 2017; Lawrence et al., 2015; Nusinow et al., 2020; Roumeliotis et al., 2017), epigenome(Barretina et al., 2012; Daemen et al., 2013; Ghandi et al., 2019; Iorio et al., 2016; Shankavaram et al., 2009) and phospho-proteome(Barretina et al., 2012; Daemen et al., 2013; Ghandi et al., 2019; Iorio et al., 2016; Marcotte et al., 2016; Shankavaram et al., 2009) across diverse panels of cancer cell lines is becoming available. Complementing these efforts, functional and phenotypic profiling of cancer cell lines using loss-of-function screens(Aguirre et al., 2016; Behan et al., 2019; Koh et al., 2012; Marcotte et al., 2016; McDonald et al., 2017; Meyers et al., 2017; Tsherniak et al., 2017; Wang et al., 2017) and small-molecule drug-response profiling has also been carried out by several laboratories(Barretina et al., 2012; Basu et al., 2013; Garnett et al., 2012; Gautam et al., 2016; Iorio et al., 2016).

Recently, the reproducibility of pre-clinical data and findings from high throughput profiling studies in cancer cell lines has been extensively investigated due to concerns of inconsistency between laboratories(Dempster et al., 2019; Gautam et al., 2019; Haibe-Kains et al., 2013; Haverty et al., 2016; Jaiswal et al., 2017; Mpindi et al., 2016; Niepel et al., 2019). In particular, the consistency of high throughput drug sensitivity screens has been questioned and re-analyzed by multiple groups(Bouhaddou et al., 2016; Geeleher et al., 2016; Haibe-Kains et al., 2013; Mpindi et al., 2016; Safikhani et al., 2016). Similarly, functional gene dependency estimates based on genome-wide RNAi screens have been reported to be relatively inconsistent, mainly due to the off-target effects inherent to the RNAi technique(Jaiswal et al., 2017). Furthermore, given the nature of cell culture techniques by which cell lines are passaged and seeded from a small population, it is likely that even identical cell lines may accumulate genomic variability and differences in their clonal composition from one research laboratory to another(Ben-David et al., 2018). This type of variability introduces an additional level of complexity which influences the repeatability of research findings and biological conclusions(Ben-David et al., 2018; Gillet et al., 2013).

In addition to the experimental issues, it is also known that the technical platform being used for high throughput measurements as well as the computational methods used in their data processing are important contributors to the consistency of research results(Haverty et al., 2016; Mpindi et al., 2016). Many of the technical platforms for molecular profiling are still in a nascent stage of development, and thus the resulting data are error prone, even when using state-of-the-art data processing and normalization procedures. Moreover, there exist major differences in the panels of cell lines profiled between research sites(Ben-David et al., 2018), hence making the comparisons and integration of profiling data intricate and biased. Therefore, there is a need for a comprehensive and quantitative analysis of the relative consistency of molecular, genetic and phenotypic characteristics of cancer cell lines from different research laboratories and technology platforms, with the aim to improve the robustness of the conclusions drawn from these studies.

In this study, we first performed a systematic statistical meta-analysis to estimate the reproducibility of various types of molecular profiles, or ‘modalities’, of cancer cell lines. Subsequently, we built on these analyses, with the aim to identify robust and reproducible gene signatures with consistent evidence across multiple research laboratories, and hence, more likely to be implicated in cancer. To do so, we developed a novel multi-omics integrative approach for jointly analysing of datasets generated from multiple studies for multiple modalities, which also accounts for differences in the panels of cell lines profiled between the research sites. Using 53 omics datasets from 12 research laboratories encompassing 9 data modalities for 2018 cancer cell lines, we demonstrate how our data driven approach is able to identify well-known driver genes of established relevance in breast cancer, as well as novel targets for therapeutic opportunities.

## Materials and Methods

### Publicly available datasets used in the multi-modal meta-analysis

#### 1) The Broad Institute, Cambridge, USA (abbreviation: BROAD)

The Broad institute is carrying out a number of large-scale cell line profiling projects such as the Cancer Cell Line Encyclopedia (CCLE)(Barretina et al., 2012; Ghandi et al., 2019), Cancer Dependency Map (DepMap)(Meyers et al., 2017; Tsherniak et al., 2017) and Cancer Therapeutic Response Portal (CTRP)(Basu et al., 2013; Seashore-Ludlow et al., 2015). Specifically, we used the point mutation profiles of coding genes among 1570 cancer cell lines from the DepMap project(DepMap, 2019). We included point mutations that were either categorized as deleterious or had FATHMM(Shihab et al., 2013) score ≤ - 0.75, and binarized the genes for presence or absence of a mutation with a functional consequence. The processed copy number profiles of 1080 cancer cell lines, generated using the Affymetrix SNP 6.0 arrays, were obtained from CCLE(DepMap, 2019). Gene-level copy number gain and losses were called using a stringent threshold of ≥ 1 and ≤ -1.5, respectively. Genome-wide transcriptomic profiles for protein-coding genes generated with RNA-sequencing for 1156 cancer cell lines were obtained from the DepMap resource(DepMap, 2019). Likewise, protein phosphorylation levels of 217 proteins in 899 cancer cell lines profiled using reverse phase protein arrays (RPPA) were obtained from the CCLE resource(Ghandi et al., 2019). Quantitative proteomic profiles for 375 cell lines were generated using TMT-labelled multiplexed protocol for sample preparation(Nusinow et al., 2020). For methylation profiles, we averaged gene-level methylation profiles of promoters situated 1kb upstream of transcription start sites for all coding genes in 843 cancer cell lines, originally generated using reduced representation bisulfite sequencing (RRBS) method). For functional profiles, we used loss-of-function data from the Achilles Project that was generated with a pooled genome-wide shRNA screening of 501 cancer cell lines(DepMap, 2019; Tsherniak et al., 2017). Gene dependency scores of each coding gene were estimated using the DEMETER2 algorithm(Tsherniak et al., 2017). We also analyzed gene dependency scores based on the genome-wide CRISPR-Cas9 knockout screens performed using various pooled sgRNA libraries from the DepMap portal(Aguirre et al., 2016; DepMap, 2019; Meyers et al., 2017; Wang et al., 2017). The Avana library was screened in 485 cancer cell lines(Meyers et al., 2017), the 120K sgRNA GeCKO v2 library was screened in 33 cancer cell lines(Aguirre et al., 2016), and the Sabatini library was screened in 15 acute myeloid leukemia (AML) cell lines(Wang et al., 2017). All the raw data for the knockout screens were processed by the Ceres algorithm(Aguirre et al., 2016) and downloaded from the DepMap or Achilles data portal. Drug response profiles of cancer cell lines estimated using the DSS2 method(Yadav et al., 2014) for CCLE and CTRP v2 were obtained from the PharmacoDB database(Smirnov et al., 2018). While CCLE screened 24 compounds against 504 cell lines, the CTRP v2 dataset was generated by screening 544 compounds against 887 cell lines. Drug sensitivity screens were based on CellTiter-Glo (CTG) assay to measure cell viabilities.

#### 2) The Sanger Institute, Hinxton, UK (abbreviation: GDSC)

The Sanger Institute has also carried out several efforts for molecular characterization of cancer cell lines, performed under the Genomics of Drug Sensitivity in Cancer (GDSC) project(Garnett et al., 2012; Iorio et al., 2016; Yang et al., 2012). We analyzed mutational profiles of 1000 cancer cell lines for coding genes from the COSMIC Cell lines project generated by whole genome sequencing(Bamford et al., 2004). We selected mutations that were categorized as pathogenic or had FATHMM score ≤ -0.75, and binarized the genes for presence or absence of a point mutation with a functional consequence. Copy number profiles for 991 cancer cell lines generated using the Affymetrix SNP 6.0 arrays and processed with the PICNIC(Greenman et al., 2010) algorithm were obtained from the GDSC portal(Yang et al., 2012). Total copy number calls made were normalized by the ploidy level as follows:

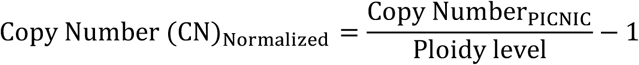

Gene copy number gains and losses were called by setting a threshold of ≥ 0.95 and ≤ -0.65, respectively, for the normalized copy number values. The RMA normalized gene expression profiles generated by Affymetrix Human Genome U219 array for 1156 cancer cell lines were available from the GDSC portal(Yang et al., 2012). For methylation profiles(Yang et al., 2012), raw intensities generated by Illumina HumanMethylation450 BeadChip were processed using the Illumina Methylation Analyzer (IMA) R package(Wang et al., 2012). Quality control was performed by removing CpG sites with missing rate >5% and detection P > 0.05. Gene level methylation intensities were obtained by averaging the methylation levels of CpG sites located in the annotated promoter site of each gene within a range of 1500 base pairs (bp) upstream of the transcription start site. Ultimately, methylation levels for ∼19000 coding genes in 1026 cancer cell lines were available for further analyses. Drug response profiles of 250 compounds tested in 1075 cell lines were quantified using the DSS2 method(Yadav et al., 2014) for GDSC1000 dataset, as obtained from the PharmacoDB database(Smirnov et al., 2018). Drug sensitivity screens were based on fluorescent nucleic acid stain probes such as Syto 60 assay to measure cell viabilities.

Additionally, proteomic and phospho-proteomic profiles of 50 colorectal cancer cell lines generated by multiplexed quantitative mass-spectrometry (MS)-based proteomics technology were made available at the Sanger Insitute(Roumeliotis et al., 2017). Multiplexing was performed by the isobaric labeling technology with tandem mass tag (TMT) reagents. Since we observed a minimal overlap between the phospho-peptides of proteins that were identified using MS in this study, when comparing to the protein residues profiled using targeted RPPA in other studies, we averaged multiple phospho-peptides corresponding to the same protein to generate protein-level phosphorylation estimates. Gene dependency profiles were generated for 324 cell lines with a pooled genome-wide CRISPR-Cas9 knockout screens performed using two pooled sgRNA libraries, the Human CRISPR Library v.1.0 and v.1.1(Behan et al., 2019). We used the copy number bias-corrected count fold changes as a measure of gene dependency in our analyses.

#### 3) Genentech Inc., USA (abbreviation: gCSI)

We used gene expression profiles and mutation calls for 675 cell lines generated by RNA sequencing(Klijn et al., 2015). We analyzed those mutations that were categorized as deleterious by variant function annotator methods: SIFT(Kumar et al., 2009), Condel(González-Pérez and López-Bigas, 2011) and PolyPhen(Ramensky, 2002). We further annotated the variants using FATHMM(Shihab et al., 2013), and selected mutations with score ≤ -0.75 and binarized the genes for presence or absence of a mutation with a functional consequence. Gene copy number profiles for 668 cancer cell lines were generated using the Illumina HumanOmni2.5 4v1 arrays and processed with the PICNIC algorithm. We used ploidy-corrected copy number calls to categorize the amplifications and deletions. Copy number gains and losses were called by setting a threshold of ≥ 0.95 and ≤ -0.65, respectively. Drug response estimates for 16 compounds tested in 409 cell lines and quantified using the DSS2 method(Yadav et al., 2014) were available from the PharmacoDB database(Smirnov et al., 2018). Drug sensitivity screens were based on CellTiter-Glo (CTG) assay to measure cell viabilities.

#### 4) National Cancer Institute, USA (abbreviation: NCI60)

The NCI-60 cancer cell line profiling data were extracted through the CellMiner data portal(Shankavaram et al., 2009), followed by further processing for the meta-analyses. Mutational profiles were generated by exome sequencing. We focused on mutations that were categorized as deleterious by the SIFT(Kumar et al., 2009) and MA(Reva et al., 2011) variant function annotators. We also annotated the variants using FATHMM(Shihab et al., 2013), and selected those variants with score ≤ -0.75, and binarized the genes for presence or absence of a mutation with a functional consequence. We used summarized log-scale intensities representing copy number profiles generated by combining probe intensities from four platforms (Agilent Human Genome CGH Microarray 44A, Nimblegen HG19 CGH 385K WG Tiling v2.0, Affymetrix GeneChip Human Mapping 500k Array Set and Illumina Human1Mv1_C BeadChip). A threshold of ≥ 0.9 and ≤ -0.9 was used to call copy number gains and losses, respectively. Similarly, processed GCRMA normalized gene expression profiles generated with Affymetrix Human Genome U133 plus 2.0 array was used. For methylation data, raw intensities generated by Illumina HumanMethylation450 BeadChip were processed to calculate gene promoter level methylation as described in the earlier section. Log intensities of protein phosphorylation site levels on 94 proteins generated using 162 antibodies by RPPA were obtained. For proteomic profiles of the NCI60 panel cell lines, we used the label-free iBAQ based quantitative estimates of protein levels(Gholami et al., 2013).

#### 5) University Health Network, Canada (abbreviation: UHN)

We downloaded the processed datasets from the Breast Functional Genomics data portal(Marcotte et al., 2016). Log ratios representing copy number profiles generated using the Human Omni-Quad BeadChip array and processed using the Circular Binary Segmentation (CBS) algorithm(Olshen et al., 2004) was available for 79 breast cancer cell lines. A threshold of ≥ 0.5 and ≤ -0.7 was used to call copy number gains and losses, respectively. Transcriptomic profiles were generated for 82 breast cancer cell lines. Log intensities of protein phosphorylation levels of 193 proteins were generated using 245 antibodies by RPPA. For gene dependency profiles, we used data from pooled genome-wide shRNA screen performed on 120 cancer cell lines from breast, pancreatic and ovarian tissue types(Koh et al., 2012; Marcotte et al., 2016). Gene dependency scores of each coding gene was estimated using the DEMTER2 algorithm(Tsherniak et al., 2017).

#### 6) Oregon Health and Science University, USA (abbreviation: OHSU_BREAST)

All the processed datasets for breast cancer cell lines were downloaded from the Synapse portal(Costello et al., 2014; Daemen et al., 2013). Transcriptomic and genomic profiles were produced by RNA sequencing and exome sequencing, respectively. Point mutations were annotated as described in earlier sections using FATHMM(Shihab et al., 2013). Log ratios representing copy number profiles generated using the Affymetrix Genome-Wide Human SNP Array 6.0 and processed with the Circular Binary Segmentation (CBS) algorithm(Olshen et al., 2004) were available for 77 breast cancer cell lines. A threshold of ≥ 0.5 and ≤ -0.7 was used to call gene copy number gains and losses, respectively. For methylation data, raw intensities generated with Illumina Infinium Human Methylation27 BeadChip Kit was used for the genome-wide detection of 27,578 CpG loci, spanning a total of 14,495 genes. Probe intensities were processed to estimate gene promoter level methylation levels as described in the earlier section. Log intensities of protein phosphorylation levels of 146 proteins generated by reverse phase protein lysate arrays were available. We utilized drug response profiles of 89 compounds tested in 71 cell lines and quantified using the DSS2 method(Yadav et al., 2014) from the PharmacoDB database (Smirnov et al., 2018). Drug sensitivity screens were based on CellTiter-Glo (CTG) assay to measure cell viabilities.

#### 7) University of Texas, MD Anderson Cancer Center, USA (abbreviation: MCLP)

We re-analyzed log-intensities of protein phosphorylation levels of 382 proteins generated using 452 antibodies by RPPA for 650 cancer cell lines were (Li et al., 2017).

#### 8) Massachusetts General Hospital Cancer Center, USA (abbreviation: MGHCC_BREAST)

Quantitative proteomic profiles of 41 breast cancer cell lines were generated using multiplexed quantitative mass-spectrometry (MS)-based proteomics technology(Lapek et al., 2017). Multiplexing was performed using the isobaric labeling technology with ten-plex tandem mass tag (TMT) reagents.

#### 9) University of Washington, USA (abbreviation: UW_TNBC)

We re-analyzed quantitative proteomic profiles of 20 breast cancer cell lines generated using non-multiplexed label-free quantitative mass-spectrometry (MS)-based proteomics technology(Lawrence et al., 2015).

#### 10) Novartis, USA (abbreviation: DRIVE)

We made use of gene dependency profiles for 8195 genes in 398 cancer cell lines for which raw data was generated by pooled genome-wide shRNA libraries(McDonald et al., 2017). shRNA level scores were collapsed to gene dependency scores of each coding gene using the DEMTER2 algorithm(Tsherniak et al., 2017).

#### 11) Institute for Molecular Medicine Finland (abbreviation: FIMM)

Drug response estimates for 52 compounds tested in 50 cell lines and quantified using the DSS2 method were obtained from the PharmacoDB database(Gautam et al., 2019; Mpindi et al., 2016; Smirnov et al., 2018). Drug sensitivity screens were based on CellTiter-Glo (CTG) assay to measure cell viabilities.

#### 12) Max Planck Institute of Biochemistry, Germany (abbreviation: MPIB_HGSOC)

We re-analyzed quantitative proteomic profiles of 30 ovarian cancer cell lines generated with a label-free quantitative mass-spectrometry-based proteomics technology(Coscia et al., 2016).

### Calculation of target addiction score

Since drug response profiles exist in the compound space, we projected them into gene space to create an additional functional data modality. To do this, we used our previously described pipeline, target addiction scoring (TAS), which transforms the drug response profiles into target addiction signatures(Jaiswal et al., 2019). The TAS pipeline makes use of drug poly-pharmacology to integrate the drug sensitivity and target selectivity profiles through systems-wide interconnection networks between drugs and their targets, including both primary protein targets as well as secondary off-targets. The TAS approach is individualized in the sense that it uses the drug sensitivity profile for each cancer cell line screened against a library of bioactive compounds, and then transforms the observed phenotypic profile into a sample-specific target addiction profile, hence enabling ranking of potential therapeutic targets based on their functional importance in the particular cell line.

We applied the TAS pipeline separately to each drug response dataset considered in the study. First, we obtained the set of potent protein targets for each drug from various drug-target databases as described previously(Jaiswal et al., 2019) (Table S1). For instance, we retrieved at least one potent target for 349/495 compounds profiled in the CTRP dataset. The rest of the compounds are either non-targeted drug treatments or compounds with unknown target profiles. Likewise, 201/250 in GDSC; 44/52 in FIMM; 33/89 in OHSU; 13/16 in gCSI; and 19/25 compounds in CCLE dataset. For each individual target *t*, TAS_t_ is calculated by averaging the observed drug response (e.g., DSS_2_) over all those *n_t_* compounds that target the protein *t.* Eventually, we were able to derive such functional TAS profiles for a median of 222 targets in each dataset:

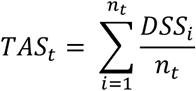

### Statistical analyses for reproducibility analyses

Spearman correlation analysis was conducted to evaluate the reproducibility of continuous molecular data types between any two studies. The reproducibility analyses were performed on the identical set of cell lines and on the common set of genes between any two datasets. Matthews correlation coefficient (MCC) was calculated to assess the consistency between binarized data types, such as gene-level mutational profiles, copy number gain and loss profiles. Only the correlation values with P < 0.05 calculated based on molecular profiles with ≥ 10 observations were considered for further analysis.

### Data processing for meta-analysis and integration

Binarized datasets of gene copy number gains and copy number losses were generated from their continuous CNV profiles using the respective thresholds, as specified above. Protein phosphorylation intensities from multiple residues mapping to the same protein were averaged to generate protein-level phosphorylation estimates. Since the GDSC phopho-proteome study in colorectal cancer cell lines used global MS technique for protein phosphorylation profiles(Roumeliotis et al., 2017), only those proteins that were also profiled in other research sites using targeted RPPA were considered for the meta-analysis.

### Meta-analysis and data integration framework

First, we define the notion of a “Cancer Cell line Specific” (CCS) gene; as a gene that exhibits a molecular attribute unique to a particular cancer cell line. In other words, a CCS gene is unique to a given cell line in reference to all the other cell lines in the particular dataset, i.e. having a context-specific property, and therefore possibly related to a specific cancer subtype or cellular function. Statistically, a gene that has the tendency to be located towards the extremes of the distribution in any given dataset is considered as a CCS gene. For instance, the expression of ERBB2 gene is much higher in HER2 driven breast cancer cell lines, compared to cell lines from other tissue types. Thus, in all HER2+ cell lines, ERBB2 is a CCS gene in the gene expression modality.

Next, we quantify the strength of evidence for cell-line specificity (CCS) for each gene across data modalities and datasets from different laboratories. We only consider data modalities in the gene space, and exclude drug-response based profiles. Specifically, we estimate the CCS evidence of a gene using its deviation from the mean over a panel of cell lines, also called as Outlier Evidence Score (OES). Further, we integrate the OES scores of genes across multiple laboratories for each data modality to identify the robust CCS (rCCS) genes, based on the rationale that, if the CCS property persists through datasets across several studies, despite the differences in the panel of cell lines that were profiled in each dataset, then the likelihood for being a robust and reproducible CCS gene increases.

For continuous data modalities, i.e. gene expression, protein expression, gene methylation, protein phosphorylation, drug sensitivity and gene dependency profiles; first an OES is estimated for each gene and specific cell line in all datasets for a given data modality across multiple laboratories. OES score is calculated as a z-score over all the cell lines profiled in that study. Then, one-class rank product analysis is performed to integrate over all OESs for each gene and cell line combination from datasets across multiple laboratories using RankProduct package(Del Carratore et al., 2017). Genes with percentage of false positives (pfp) below a pre-specified threshold were considered statistically significant and defined as CCS genes for the respective data modality. For methylation, gene expression, protein expression and protein phosphorylation data modalities, genes with pfp < 0.10 were considered significant. For functional gene dependency profiles, genes with pfp < 0.2 were considered significant owing to the higher technical variability in these datasets.

For cell lines that were profiled in only one study, the rank product analysis could not be performed, and therefore all the top-ranked genes with OES below a specified threshold were selected: top-100 genes (0.5% of all genes) were selected for all continuous variable datasets; except for functional datasets (Achilles, DepMap, DRIVE and UHN), the top-200 genes were considered (1% of all genes). For the protein phosphorylation datasets, the top-10 genes (0.5% of all proteins) were selected, considering that 200 proteins were assayed by reverse phase protein arrays on average.

For binarized data modalities, i.e. copy number gain and loss and gene point mutation profiles, we defined a Proportion Score (PS), which is the proportion of cell lines in which the particular genomic alteration is observed. Next, for a given cell line, all PSs for that data modality type were combined for each individual gene from different datasets. Finally, any alteration that has been observed in ≤ 10% cell lines (arbitrary selected threshold) in any single dataset, is considered as a CCS gene.

Genes that are identified as CCS genes in two or more data modality types are considered in our analyses as robust CCS (rCCS) genes. Fisher’s exact test was performed to evaluate the difference in the proportion of rCCS genes between different cancer subtypes or pre-defined subgroups of cancer cell lines. Overall, we applied the CLIP to 1047 cancer cell lines for which data was available for ≥ 6 modalities (Table S2).

### Genetic interaction and synthetic lethal analyses

To identify genetic interactions between genes, Fisher’s exact test was performed to evaluate the difference in the proportion of rCCS genes between two groups of cancer cell lines, mutated and wild-type, for all the well-known cancer driver genes. Out of the 2018 cell lines that were included in the meta-analyses, we considered 1047 cell lines for which molecular data were available in ≥ 6 of the 8 data modalities in gene-space. Further, we removed cell lines derived from bone, skin, nervous and hematopoietic systems, restricting our analyses to epithelial cancer cell lines (n=679). For selecting mutated driver genes with relevance in patient tumors, we considered the highly frequent driver genes in patient tumors from a recent pan-cancer study of mutational landscape in The Cancer Genome Atlas (TCGA) dataset(Kandoth et al., 2013), and a subset of the driver genes that were mutated in at least 5 cell lines. For defining the set of potential synthetic lethal (SL) interactions, we considered only those rCCS genes that were also identified as essential genes, based on the evidence from the gene dependency modality (Table S3).

### Survival analysis in the patient tumor cohorts

Kaplan–Meier survival analysis and univariate Cox proportional hazard test was used to assess the difference in overall survival between any two pre-defined patient groups from the TCGA pan-cancer dataset; the sub-groups were defined based on cell line predictions, either on the basis of methylation levels of ECHDC1 in breast cancer tumors or expression levels of DDX27 in endometrial, liver and kidney tumors.

### Experimental validation of ECHDC1 knockdown

Noncancerous epithelial cell line MCF10A and breast carcinoma cell line BT-474 (both American Type Culture Collection; ATCC) were cultured according to manufactureŕs instructions in a humidified incubator with 5% CO_2_ at 37 °C and routinely checked and tested negative for mycoplasma contamination using MycoAlertPlus^TM^ Mycoplasma Detection Kit (Lonza) according to manufacturer’s instructions. Small interfering RNA (siRNA) reagent against human ECHDC1 (GEHealthcare, Dharmacon SMARTPool: L-009522-01-00005) and non-silencing control siRNA (GEHealthcare, Dharmacon: D-001206-14-20) were transfected using Lipofectamine 2000 (Thermo Scientific) according to manufactureŕs protocol.

Cells were transfected on 2D and cultured for 8 hours before embedding as single cells in 3D hydrogels composed of collagen or Matrigel. For collagen matrix, rat tail collagen type I (Sigma-Aldrich) was dissolved in 0.25% acetic acid and diluted 1:1 with 2 x MEM (Gibco) to final concentration of 2.25 mg/ml. Matrigel (Corning, 354263) was diluted to 12 mg/ml with HBSS (Sigma-Aldrich). Cells (3 x 10^3^ cells /ml) with transient knockdown were embedded in 3D collagen and Matrigel and cultured in complete media. Cell invasive growth was followed for 5 days by phase-contrast imaging using inverted epifluorescence microscope (Axiovert 200; Carl Zeiss).

### Metabolite assay of intermediates in ECHDC1 pathway

A total of 14 breast cell lines (see Table S4), 7 for ECHDC1 rCCS positive (rCCS+) and 7 for rCCS negative (rCCS-), were selected for sub-sequent metabolite assays based on their ECHDC rCCS status. In total, 42 samples (14 lines with 3 replicates) were run with UPLC-MS (MRM) method, each sample containing ca. 4 x 10^-06^ cells. Culture conditions for each cell line are detailed in Table S4.

Quenching protocol for adherent cells involved the following steps: cells were washed with 2x volume of cold phosphate-buffered saline (PBS) and incubated with trypsin (TrypLE™ Express Enzyme (1X), #12605010, Invitrogen) at 37 °C until the cells detached. Trypsin was inactivated by adding an equal volume of cold fetal bovine serum (FBS, #10270-106, Invitrogen). The cells were counted and centrifuged at 400 x g for 5 minutes at 4 °C. The cells were washed with 2x volume of cold PBS, each time centrifuging at 400 x g for 5 minutes at 4 °C. For each sample 4 x 10^-06^ cells were resuspended in 500 μl of cold PBS and centrifuged at 400 x g for 5 minutes at 4 °C in 1.5 ml microcentrifuge tube. The cells were quickly washed with 2x volume of deionized water, not disturbing the pellet and not exposing the cells to water for more than 4-5 seconds. All water was aspirated from the tube. The cells were frozen in liquid nitrogen and stored at -80 °C.

The protocol for cell disruption and extraction was adapted from a previously described method(Dettmer et al., 2011). Briefly, cells disrupted with a combination of freeze-thaw cycle (−80/+4°C) and sonication. 1000µl of 80% MeOH (Honeywell, Riedel-de-Haën™, Seelze. Germany) and 10µl of internal standard (ISTD) (conc. 10 µg/ml) was added to the purified cells. Then samples were ultra-sonicated in ice bath for 10 minutes, put to liquid nitrogen back and forth three times. After cell disruption, samples were vortexed for an hour, centrifuged at 21 500 g for 5 minutes at +4 °C. The second extraction was performed with 600 µl of 100% MeOH for 30min. The supernatant (1600µl) was dried under vacuum (MiVac Duo concentrator, GeneVac Ltd, Ipswich, UK), reconstituted to 50µl of 0.1% formic acid in acetonitrile (ACN)/H_2_O (Honeywell, Riedel-de-Haën™, Seelze. Germany) and run with UPLC-QTRAP/MS with ESI (+) and (-) switching (ExionLC UPLC, ABSciex; 6500+ QTRAP-MS, ABSciex).

Ten microliter (μl) of extract was injected into the LC column with the mobile phase flow of 0.4 ml/ minute at +35°C. The LC separation was carried out on a reversed-phase UPLC-column (Waters Acquity BEH C18, 150 × 2.1 mm, Ø 1.7µm). A gradient elution of the analytes was achieved using a program with mobile phases A (aqueous 0.1% formic acid) and B (0.1% formic acid in ACN). The linear gradient started at 99% A and 1% B, was held for 2 minutes and proceeded from 1% B to 10% in 2 minutes, 10%B to 90% in 2 minutes, held at 90% for 1min, then switched back to 1% B and left to stabilize for 2 minutes. Total run time of 9 minutes.

In total, 12 metabolites were analyzed from the cells; aspartate (Asp), glutamate (Glu), glutamine (Gln), α-ketoglutarate (α-KG), fumarate, succinate, malate, citrate, isocitrate, 2-hydroxy-3-methylbutyrate (2OH-3MBA), malonate, methylmalonate (MMA). D_3_-methylmalonate (D_3_-MMA) was used as an ISTD. Multiple Reaction Monitoring (MRM) transitions for 13 analytes were monitored: m/z 134/74 for Asp, 148/56 for Glu, and 147/84 for Gln in positive mode; and m/z 145/83 for α-KG, 114.9/71 for fumarate, 117.1/73 for succinate, 132.8/71 for malate, 190.8/111 for citrate, 190.8/73 for isocitrate, 117.1/7 for 2OH-3MBA, 103/59 for malonate, 117/73 for MMA and 120/76 for D_3_-MMA (ISTD) in negative mode. Calibration curves with five standard mixes (conc. 100, 10, 1, 0.1 and 0.01 μg/ml) were used for quantification for all 12 metabolites. The samples were not normalized to cell count, but only for ISTD, hence unit μg/ml.

### Prediction of ECHDC1 function with gene co-regulation analysis

We used the Gene-Module Association Determination (G-MAD) algorithm, implemented in the GeneBridge toolkit(Li et al., 2019), to predict the gene function of ECHDC1. G-MAD considers transcriptome data sets from six species (human, mouse, rat, fly, worm, and yeast), and performs a competitive gene set testing method—Correlation Adjusted MEan RAnk gene set test (CAMERA)(Wu and Smyth, 2012), which adjusts for inter-gene correlations to compute the enrichment between gene-of-interest and biological modules. Gene-module connections with enrichment *P*-values that survive multiple testing corrections of the gene or module numbers are scored from 1 or −1. We restricted our analysis to breast tissue datasets (n=153) compiled in the GeneBridge expression database to enrich for tissue specific associations (Table S5).

## Results

### Compilation of available molecular modalities of cancer cell lines

We processed, re-analyzed and harmonized curated datasets of various data modalities for cancer cell lines that were generated at 12 research sites (see Material and Methods). We focused on analyzing data modalities that are available as quantitative measurements for various attributes of protein-coding genes, including methylation, mutational status, copy number alteration status, gene and protein expression, and protein phosphorylation. We further considered functional profiles such as gene-dependency estimates from loss-of-function screens and drug-response measurements (Figure 1A), and calculated an additional functional data modality by transforming drug response profiles to target protein addiction signatures. Overall, a given cell line can maximally have data across 9 modalities generated at one of the research laboratory sites (Figure 1B). The number of cell lines profiled for each data modality ranged from 171 (protein expression) to 1689 (mutation profiles, Figure 1C), making the data integration challenging for the meta-analysis (Table S6).

**Figure 1:**
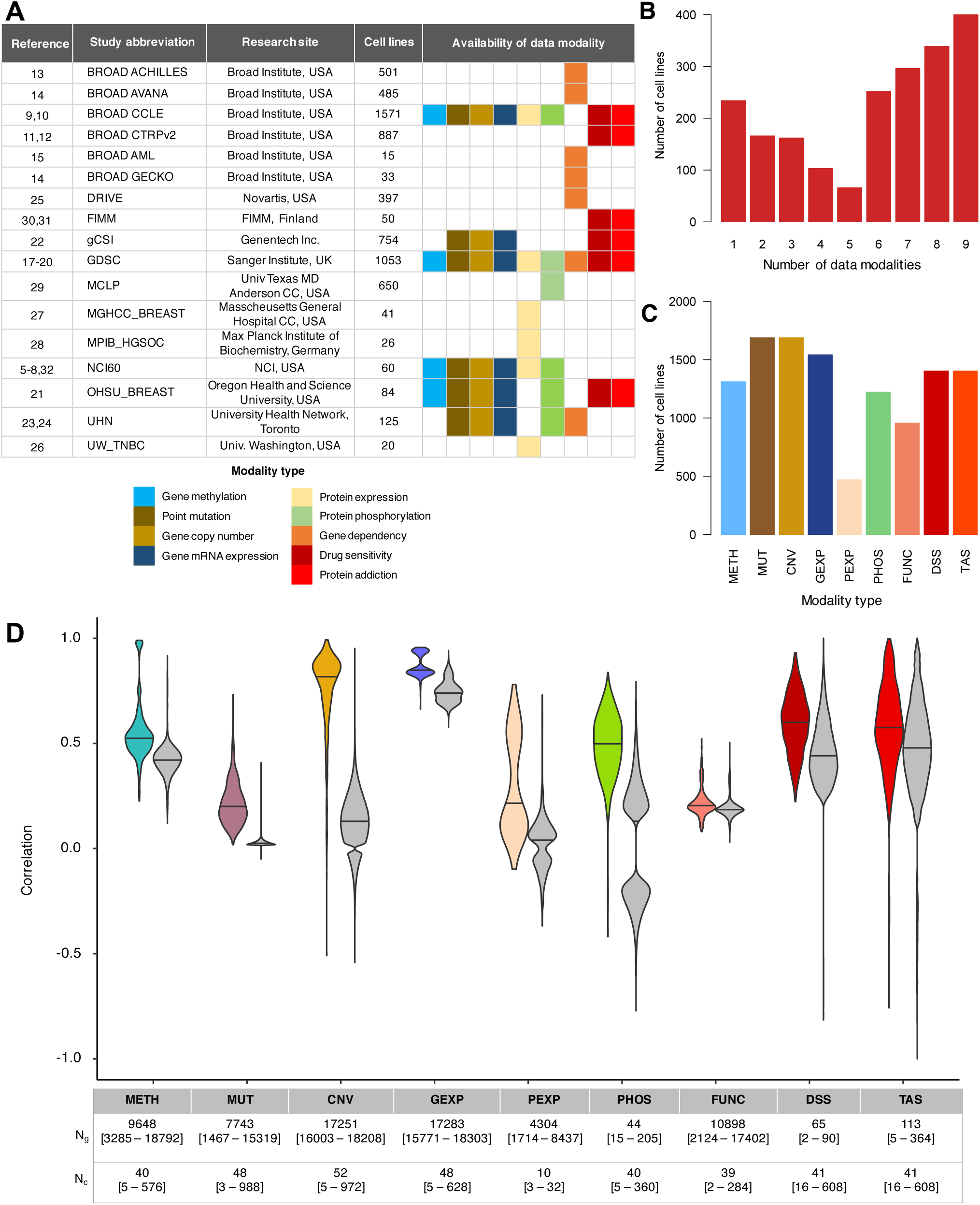
Overview of data modalities and their consistency. (A) Overview of datasets, research sites, and molecular modalities that were analyzed in the study. (B) The number of cell lines having data for the 9 types of modalities that were analyzed in the study. (C) The number of cell lines for which data were available for each of the modality types. (D) Correlation of the different types of data modalities of cancer cell lines profiled at multiple research sites. Spearman correlation was calculated between identical cell lines for the shared set of genes (i.e., the dimension of the profile) that were overlapping between any two datasets. Gray distributions show the correlation of non-identical cell lines between datasets from various research sites for comparison. N_g_ and N_c_ indicate the median [ranges] of the number of genes and cell lines, respectively, across the pairwise comparisons made between datasets from different research sites. More details on the breakdown of N_c_ and N_g_ by data modality and research site is available in Figure S1B and S2, respectively, and Figure S3 shows the correlation P-values adjusted for the sample size (N_g_). For the point mutation view, only those genes having mutations with an associated functional consequence were considered in the Matthews correlation analysis. Only those datasets for which the mutation profiles were obtained using the whole-exome sequencing technology were considered in this study.

For instance, the National Cancer Institute (NCI) programme (NCI-60) has extensively characterized a panel of 60 cancer cell lines representing 9 different cancer types(Shankavaram et al., 2009). In contrast, more large-scale efforts such as the Genomics of Drug Sensitivity in Cancer (GDSC)(Behan et al., 2019; Garnett et al., 2012; Iorio et al., 2016; Roumeliotis et al., 2017; Yang et al., 2012), Cancer Cell Line Encyclopedia (CCLE)(Barretina et al., 2012; Basu et al., 2013; Ghandi et al., 2019; Meyers et al., 2017; Seashore-Ludlow et al., 2015; Tsherniak et al., 2017), and the Genentech Cell Screening Initiative (gCSI)(Klijn et al., 2015) have characterized approximately 1000, 1500 and 675 cancer cell lines, respectively, representing a wide variety of cancer types. These studies have also performed phenotypic profiling of drug sensitivity against a library of small molecules (Figure 1A). Likewise, the DepMap project has systematically characterized the functional-genomic landscape of ∼500 cancer cell lines using genome-wide RNAi screens and several versions of genome-wide CRISPR-Cas9 loss-of-function screening libraries(Aguirre et al., 2016; Meyers et al., 2017; Tsherniak et al., 2017; Wang et al., 2017) (Figure 1A). Protein phosphorylation levels using reverse-phase protein arrays (RPPA) and phospho-proteomics have also been generated at the MD Anderson Cell Lines Project (MCLP) for 340 unique cancer signaling related proteins in ∼650 cancer cell lines(Li et al., 2017); at CCLE for 174 proteins in ∼900 cell lines (Figure S1-2).

Complementing these large-scale pan-cancer programmes, we also re-analyzed datasets from more targeted efforts that have profiled cell lines of a specific cancer type; these smaller-scale studies were included in this meta-analysis to increase the information content on selected tissue lineages. Specifically, multi-modal datasets were generated at the University Health Network (UHN)(Koh et al., 2012; Marcotte et al., 2016) at Toronto and the Oregon Health and Science University (OHSU)(Costello et al., 2014; Daemen et al., 2013) studies for >80 breast cancer cell lines. Further, proteome-scale expression levels in several breast, ovarian and colorectal cancer cell lines have been generated using mass spectrometry (MS) at the University of Washington (UW_TNBC)(Lawrence et al., 2015), Massachusetts General Hospital Cancer Center (MGHCC_BREAST)(Lapek et al., 2017), Max Planck Institute of Biochemistry (MPIB_HGSOC)(Coscia et al., 2016) and the Sanger Institute (GDSC)(Roumeliotis et al., 2017); however, the total number of cell lines profiled for MS protein expression was only 171. Further, an in-house drug sensitivity profiling dataset of >50 pan-cancer cell lines generated at the Institute for Molecular Medicine Finland (FIMM) was also utilized in the study(Gautam et al., 2019; Mpindi et al., 2016; Smirnov et al., 2018).

To enable the meta-analysis between studies, we only considered datasets that were generated in a sufficiently larger panel of cell lines (n >10), and therefore excluded datasets below the threshold. In statistical analyses, we assumed that the same cell lines profiled at each site were cultured independently. All together, we processed and re-analyzed 53 datasets, encompassing 9 modalities generated at the 12 research study sites. In total, we analyzed data for 2018 cancer cell lines having measurements for at least one of the data modalities. A substantial proportion of cell lines (>1047) had data available for ≥ 6 modalities, thus serving as a comprehensive resource for further analyses (Figure 1 B, C). Even though most cell lines had data available from multiple sites, there were ∼215 cell lines that had data available from only one study site (Figure S1 A-B, Figure S2). We reasoned that the substantial overlap between cell lines across multiple molecular layers between more than two sites provides a solid basis to perform a quantitative assessment of the reproducibility of the multiple modalities of cancer cell lines.

### Reproducibility of molecular modalities of cancer cell lines from multiple sites

We performed a systematic correlation analysis to evaluate the consistency of gene level quantitative measurements of the various data modalities from identical cell lines profiled across different research sites. Overall, we observed a wide variation in the degree of pairwise agreement between the research laboratories (Figure 1D, Figure S3). Consistent with previous observations(Haibe-Kains et al., 2013; Haverty et al., 2016; Klijn et al., 2015), copy number variation (CNV) profiles and transcriptomic profiles of cell lines were highly correlated between different study sites (Spearman correlation r_CNV_ = 0.76 [-0.51-0.99] and r_GEXP_ = 0.87 [0.66-0.96]) (Figure 1D). However, we observed a considerable range of variation in the pairwise correlation of CNV profiles between different sites, suggesting that the cell lines with poor agreement may have substantially been diverged and undergone clonal selection during cell culture. On the contrary, transcriptomic profiles tended to be highly correlated and did not exhibit such a high dispersion. Surprisingly, mutational profiles of genes harboring point mutations with functional consequences were less consistent between various study sites (r^2^_MUT_ = 0.22 [0.02 - 0.73]).

In general, methylation profiles of cell lines, corresponding to methylation levels of CpG sites located at transcription start sites of genes, were moderately consistent (r_METH_ = 0.56 [0.23 - 0.99]) (Figure 1D), and also exhibited a relatively wide range of correlation values between study sites. Likewise, protein level phosphorylation profiles were only modestly reproducible between different sites, suggesting that the targeted reverse phase protein array (RPPA) technique is relatively noisy (r_PHOS_ = 0.49 [-0.42 - 0.84]). The correlation of the global proteome expression profiled with MS was even lower, on average, and it also exhibited a wide range of variability in the relatively small number of available breast and ovarian cancer cell lines (r_PEXP_ = 0.29 [-0.09 - 0.78]). However, when considering the dimension of the profiles (median of 44 for PHOS, and 4304 for PEXP), the global protein expression correlations had higher significance on average (Figure S3). As observed previously(Haverty et al., 2016; Mpindi et al., 2016), we also found that the reproducibility of drug sensitivity profiles between sites were moderately high (r_DSS_ = 0.63 [0.22 - 0.95]), similar to the reproducibility of TAS profiles (r_TAS_ = 0.56 [-0.75 – 0.99]). In contrast, gene dependency estimates based on loss-of-function RNAi and CRISPR screens exhibited rather poor reproducibility (r_FUNC_ = 0.21 [0.08 - 0.52]).

Given that the distributions of data modalities are quite different, the correlation estimates (either Spearman or Matthew’s coefficient) are not directly comparable. To set a reference point for the pairwise comparisons, we further estimated the correlation of non-identical cell lines between the different studies (Figure 1D, grey distributions). This analysis is also useful for assessing the expected baseline correlation of different modality types. As expected, the average correlation of mRNA expression profiles of non-identical cell lines was generally high (r_GEXP_ = 0.75), suggesting that the transcriptomic landscapes are quite similar across cancer cell lines. Compared to the average correlation of non-identical cell lines, we observed a 1.17-fold increase in the mean correlation of the identical cell lines for gene expression profiles (P < 10^-10^, Wilcoxon test). We observed a similar fold increase for methylation (1.33-fold, P < 10^-10^), gene dependency (1.13-fold, P < 10^-10^) and drug response profiles (1.59-fold, P < 10^-10^). In contrast, a much higher fold increase in the correlation of identical cell lines was observed for CNV (5.9-fold, P < 10^-10^), point mutation (7.8-fold, P < 1.0×10^-10^), protein phosphorylation (27.9-fold, P <10^-10^) and protein expression profiles (12.2-fold, P < 1.5×10^-08^).

### Reproducibility of technical platforms used to generate the data modalities

Correlation analysis of the various data modalities implied the existence of bi-modal distribution of consistency estimates for some of the data modalities (Figure 1D, Figure S4). We therefore further stratified the correlation analyses separately for each of the experimental techniques to investigate whether the platform used to generate the data explains the observed variability. For instance, we observed a significantly higher reproducibility of methylation profiles between studies generated using the Illumina 450K BeadChip, compared to the correlation of methylation profiles of datasets generated using different platforms, one using array-based Illumina 450K BeadChip and the other using bisulfite sequencing-based methylation profiles (r_METH_ = 0.97 for 450K/450K vs. r_METH_ = 0.51 for 450K/Bisulfite, P < 1.0×10^-10^, Wilcoxon test) (Figure S4 J). Likewise, methylation profiles generated with the Illumina 27K BeadChip were more correlated with the Illumina 450K profiles than with bisulfite sequencing-based profiles (r_METH_ = 0.72 for 27K/450K vs. r_METH_ = 0.45 for 27K /Bisulfite, P < 1.0×10^-10^).

As expected, we observed a significantly better agreement between those studies in which the transcriptomic profiles of cell lines were measured using RNA sequencing, as compared to the microarray-based technologies (r_GEXP_ = 0.93 for RNA-seq vs. r_GEXP_ = 0.84 for arrays, P < 1.0×10^-10^, Wilcoxon test) (Figure S4). Interestingly, moderate differences were also observed for the reproducibility of protein phosphorylation profiles generated with RPPA and MS technologies. Specifically, RPPA-based protein phosphorylation profiles were slightly better correlated with studies based on RPPA than with MS (r_PHOS_ = 0.45 for MS/RPPA vs. r_PHOS_ = 0.49 for RPPA/RPPA, P = 0.03) (Figure S4 K). Moreover, we observed that drug sensitivity screens, and TAS profiles based on CellTiter-Glo (CTG) assay were significantly more correlated in comparison to those based on fluorescent nucleic acid stain probes such as Syto 60 (r_DSS_ = 0.72 for CTG/CTG vs. r_DSS_ = 0.55 for CTG/Syto60, P < 10^-10^) (Figure S4 L).

In the comparison of gene dependency profiles obtained either from genome-wide RNAi knock-down or CRISPR knockout screening techniques (Figure 2A), we observed a relatively low correlation between functional studies based on genome-wide RNAi screens (r_FUNC-RNAi_ = 0.22) (Figure 2B), in line with previous reports showing that gene dependency profiles based on this technique are less robust(Gautam et al., 2019; Jaiswal et al., 2017). In contrast, genome-wide CRISPR screens exhibited a moderate consistency between studies (r_FUNC-CRISPR_ = 0.36), significantly higher compared to genome-wide RNAi screens (P < 10^-10^). As reported before(Gautam et al., 2019), the correlation between studies based on RNAi and CRISPR screens was also quite poor (r_FUNC-RNAi/CRISPR_ = 0.19) (Figure 2A). Moreover, the agreement between the two screens performed at GDSC and BROAD Avana library was slightly lower compared to the screens performed exclusively at BROAD (r_FUNC_ = 0.35 for BROAD-Avana/GDSC vs. r_FUNC_ = 0.43 for BROAD-Avana/GeCKO/AML, P = 2.7×10^-08^). These results demonstrate how laboratory-specific factors contribute to differences in the quantitative estimates of gene-dependency profiles.

**Figure 2:**
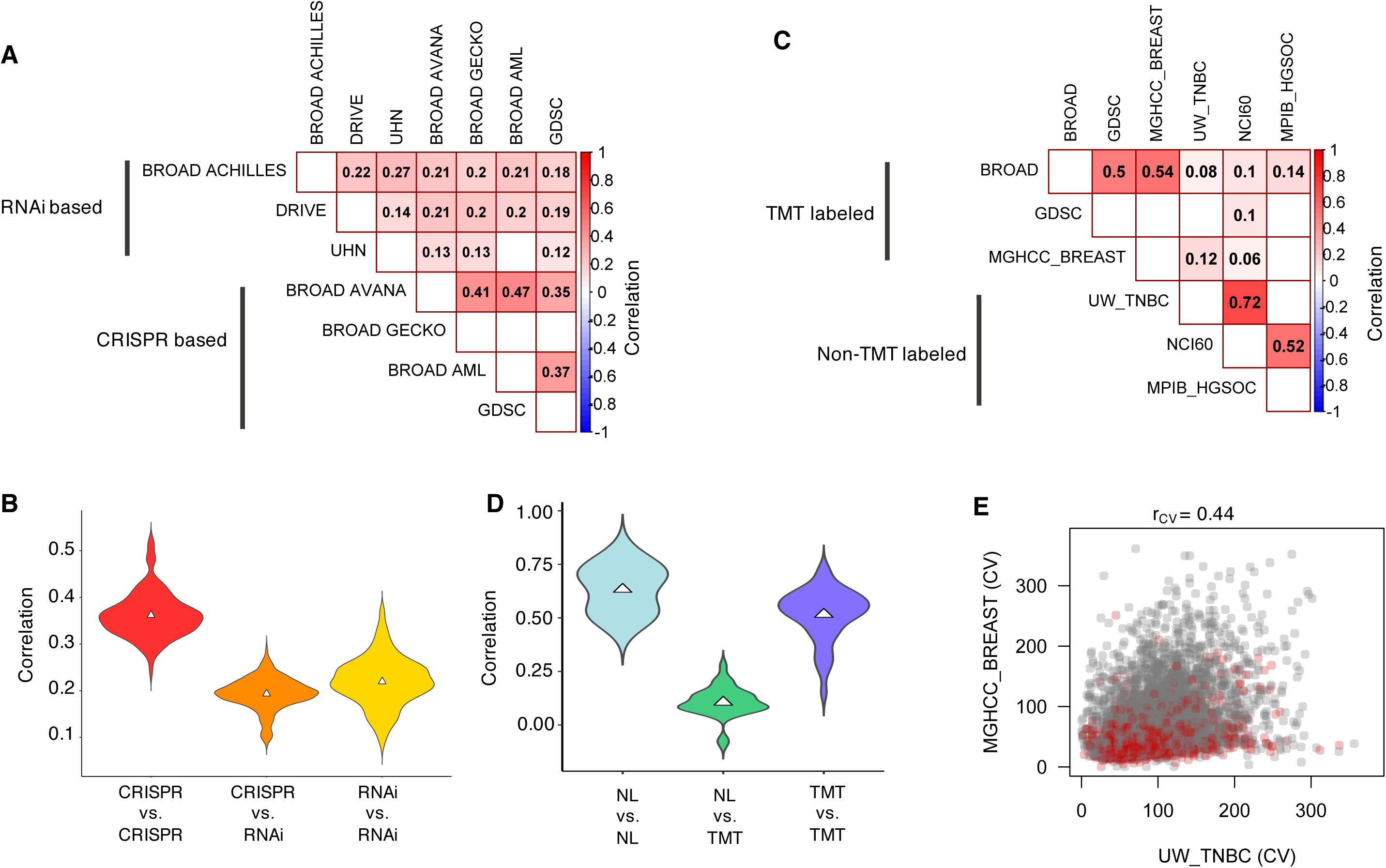
Contribution of lab-specific factors to the reproducibility of functional gene dependency profiles and MS-based proteomic profiles. (A) Average Spearman correlation of gene dependency profiles of cancer cell lines calculated based on genome-wide RNAi screens and CRISPR screens. The empty cells indicate that no identical cell lines were profiled between the two datasets. (B) Distribution of Spearman correlation of gene dependency profiles between different study sites. (C) Average Spearman correlation of MS-based proteomic profiles between different study sites generated using different peptide labeling procedures. The empty cells indicate that no identical cell lines were profiled between the two datasets. (D) Distribution of Spearman correlation of MS-based proteomic profiles between different study sites. (E) Coefficient of variation (CV) of proteins detected and quantified in UW TNBC study (non-TMT labeled) vs. MGHCC BREAST study (TMT-labeled). Housekeeping genes are highlighted as red dots. Spearman correlation (r_cv_) was calculated to estimate the agreement in the CV estimates of common set of proteins between the two studies.

When investigating potential reason for the bi-modal distribution of correlation estimates for the MS-proteomic datasets, we found that the agreement of protein expression profiles varied depending on the sample preparation method (Figure 2C), Specifically, the BROAD, MGHCC_BREAST and GDSC studies utilized tandem mass tag (TMT)-based peptide labelling before protein abundance quantification, whereas the other studies used a non-labeled (NL) approach. The correlation between TMT-labeled and NL proteome profiles was poor (r_PEXP_ = 0.11), compared to proteome profiles generated at different study sites using the same method (r_PEXP_ = 0.63 for NL/NL and r_PEXP_ = 0.52 for TMT/TMT) (Figure 2C, D). The NL proteomic profiles were more reproducible in comparison to TMT labelled profiles (P = 0.045, Wilcoxon test). However, we observed a slightly better agreement in the coefficient of variation (CV) calculated for the common set of proteins between one TMT and another NL study (r_CV-PEXP_ = 0.44, Figure 2E). This suggests that the labelling procedure has a drastic impact on the quantitation of protein abundance, which may lead to variability in the proteomic profiles.

### An analytical framework for meta-analysis and integration of multi-modal datasets

The availability of multiple data modalities of molecular profiles of cancer cell lines from multiple studies and laboratories, that show only a moderate overlap and consistency, pose a challenge for integrative approaches that leverage the multiple levels of profiling information to identify robust driver genes and biological processes. We hypothesized that genes that have a consistent molecular pattern shared across multiple studies and modalities are more likely to have a functional consequence relevant for cancer. Towards this end, we developed a non-parametric, rank-based framework, named Cell Line specific gene Identification Pipeline (CLIP), which enables systematic meta-analysis and integration of all the datasets collected and processed in this study (Figure 3). To boost the statistical power toward finding robust and reproducible signals in these data, the CLIP framework accounts for the substantial variability in the consistency of the various types of modalities between laboratories. To overcome the problem of data sparsity we developed an individual cell line specific “bottom-up” approach of meta-analysis.

**Figure 3:**
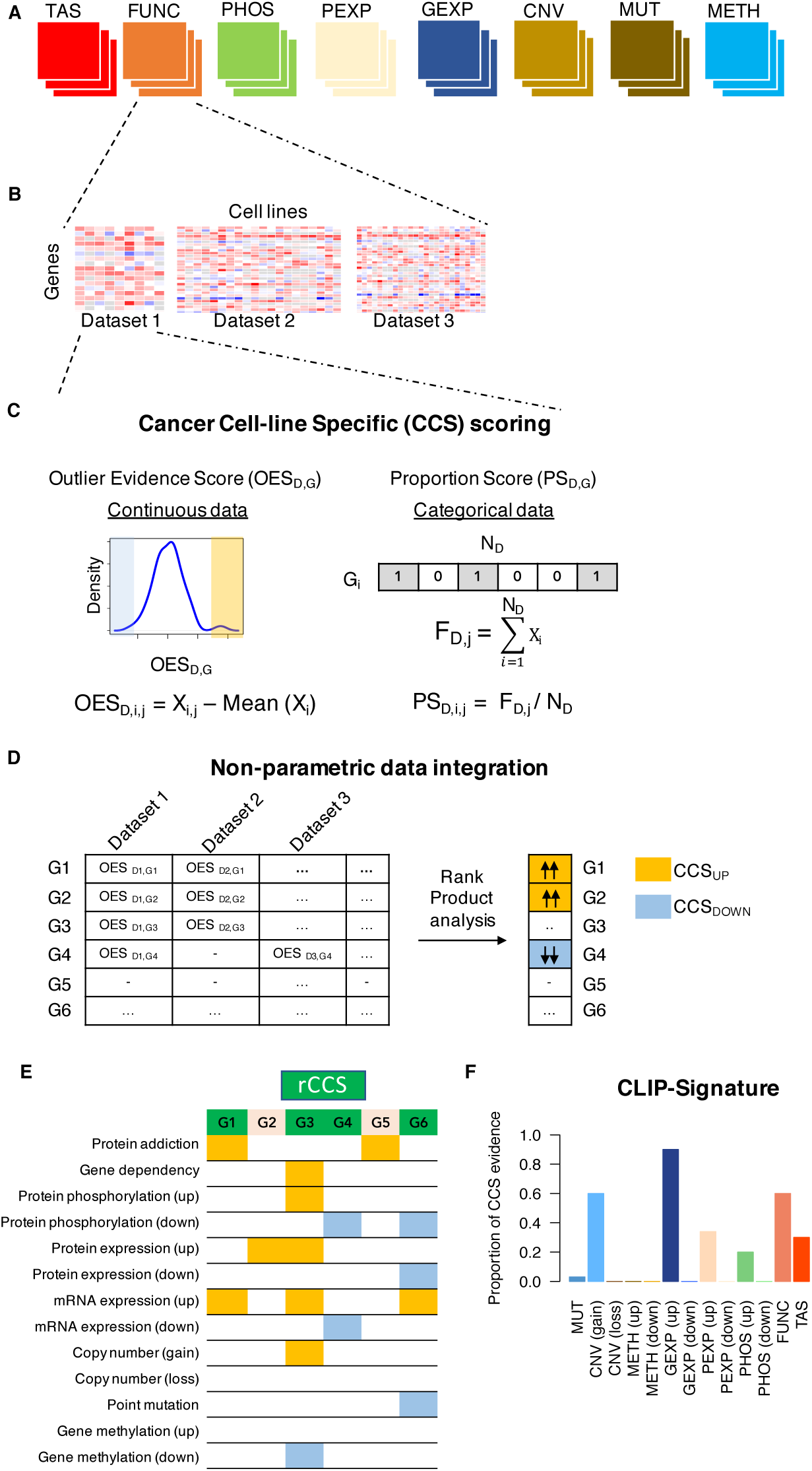
Overview of the Cell Line specific gene Identification Pipeline (CLIP) for integration of molecular datasets from multiple studies. (A) CLIP performs a meta-analysis of datasets from multiple sites for each data modality type: Target addiction score (TAS), Gene dependency (FUNC), protein phosphorylation (PHOS), protein expression (PEXP), gene expression (GEXP), copy number variation (CNV), point mutation (MUT) and methylation (METH) profiles. (B) For each modality type, CLIP iterates over datasets available from multiple sites and quantifies the cancer context specificity (CCS) property for every gene G in cell line j. (C) For all unique cell lines, the CSS property is quantified for each gene G in a dataset D. For continuous modalities (METH, GEXP, PEXP, PHOS, FUNC, TAS), we defined the Outlier Evidence Score (OESG,D,j), calculated by normalizing the observed value by the mean in the dataset for each gene (Xi). For binary modalities (CNV-GAIN, CNV-LOSS and MUT), we defined the Proportion Score (PSG,D,j) for each gene G in cell line j, calculated as the frequency of the alteration (FD,j) normalized by the total samples in each dataset (ND,j). (D) For a given cell line j, OESG,D scores across the available datasets are integrated using the Rank Product analysis to find statistically consistent genes that are either up-regulated (CCSUP) or down-regulated (CCSDOWN). (E) Finally, CLIP produces a profile of all the genes that are identified as CCS. In total, 13 different modality features were assessed by the CLIP framework, provided there are data available for a cell line for all the molecular datatypes. All genes identified as a CCS gene in any modality are highlighted, light orange for up-regulation and light blue for down-regulation. Genes that have CCS evidence across two or more modality types are considered in our analyses as robust Cancer Context Specific (rCCS) genes, highlighted as light green. (F) A schematics of CLIP signature of a gene, which summarizes its CCS evidence in a selected subset of cell lines, defined as a group based on any relevant criteria (the example shows all HER2+ breast cancer cell lines). Y-axis is the ratio of number of cell lines in which the gene is identified as a CCS gene vs. the total number of cell lines in the particular subset.

To solve the data sparsity challenge, we developed a ‘bottom-up’ meta-analysis approach based on the concept of cancer cell-line specific (CCS) genes. A CCS gene exhibits a molecular feature that is unique for a given cell line in reference to the other cell lines, i.e., CCS gene has a context-specific property, which may potentially contribute to the unique biological characteristics of the particular cell line. Statistically, CCS genes have the tendency to be located towards the extremes of a data modality distribution. For instance, the expression of ERBB2 gene is much higher in ERBB2 (HER2) driven breast cancer cell lines, compared to cell lines from other tissue types (Figure S5). The strength of a CCS property for each gene in a dataset was quantified using its deviation from the mean, as an outlier-based scoring metric (see Methods). Moreover, the statistical power and strength of quantitative evidence for a CCS gene comes from the total number of other cell lines against which the expression level of the candidate CCS gene is being compared. Here, we quantified the CCS property of each gene across all 8 types of data modalities (Figure 3A, B).

More specifically, we quantified the CCS strength of each gene in a given cell line and in each individual dataset using an Outlier Evidence Score (OES) for continuous data modalities and Proportion score (PS) for binary modalities (Figure 3C). Next, for any given cell line, OES and PS scores of all the coding genes for a specific data modality from different studies were integrated to identify those genes that were consistently “up” or “down”-regulated in a given cell line, CCS_UP_ and CCS_DOWN_ (Figure 3D). Conceptually, the two categories define a particular property of a gene, for instance, gene expression level higher or lower in the particular cell line compared to all the other cell lines. Ultimately, for each cell line, the CLIP meta-analysis framework provides a list of genes that show statistically robust evidence for being CCS genes by considering all the 8 types of molecular modalities in gene-space, and generates a cell line specific CLIP-Signature for each gene (Figure 3E). The CLIP-Signature can be used to select genes that have the strongest CCS evidence across two or more modalities, termed as robust Cancer Context Specific (rCCS) genes (Figure 3F).

### CLIP identifies established breast cancer cell line and subtype-specific drivers

To systematically test our meta-analysis approach, we reasoned that the list of rCCS genes for each cell line should be enriched for genes that determine the unique phenotypic or molecular characteristics of the particular cell line. As a proof-of-principle, we applied the CLIP framework specifically to 103 breast cancer cell lines (Figure S6), as they have been extensively profiled by multiple studies. Reassuringly, the meta-analysis approach was able to identify previously-established driver kinases in several breast cancer cell lines(Szwajda et al., 2015): ERBB2 in BT474, SKBR3 and MDAMB453, BRAF in DU4475 and PIK3CA in T47D, MCF7 and MDAMB453 cell lines (Figure 4A). The CLIP-Signature further revealed that most of the driver genes were selected based on the target addiction and gene dependency modalities, and some others based on protein-phosphorylation (up) and gene copy number (gain), as well as based on their point mutation views. Thus, in addition to identifying known drivers in breast cancer cell lines, the CLIP-Signature also provided insights into the mechanistic basis of the drivers based on multiple levels of supporting evidence across data modalities.

**Figure 4:**
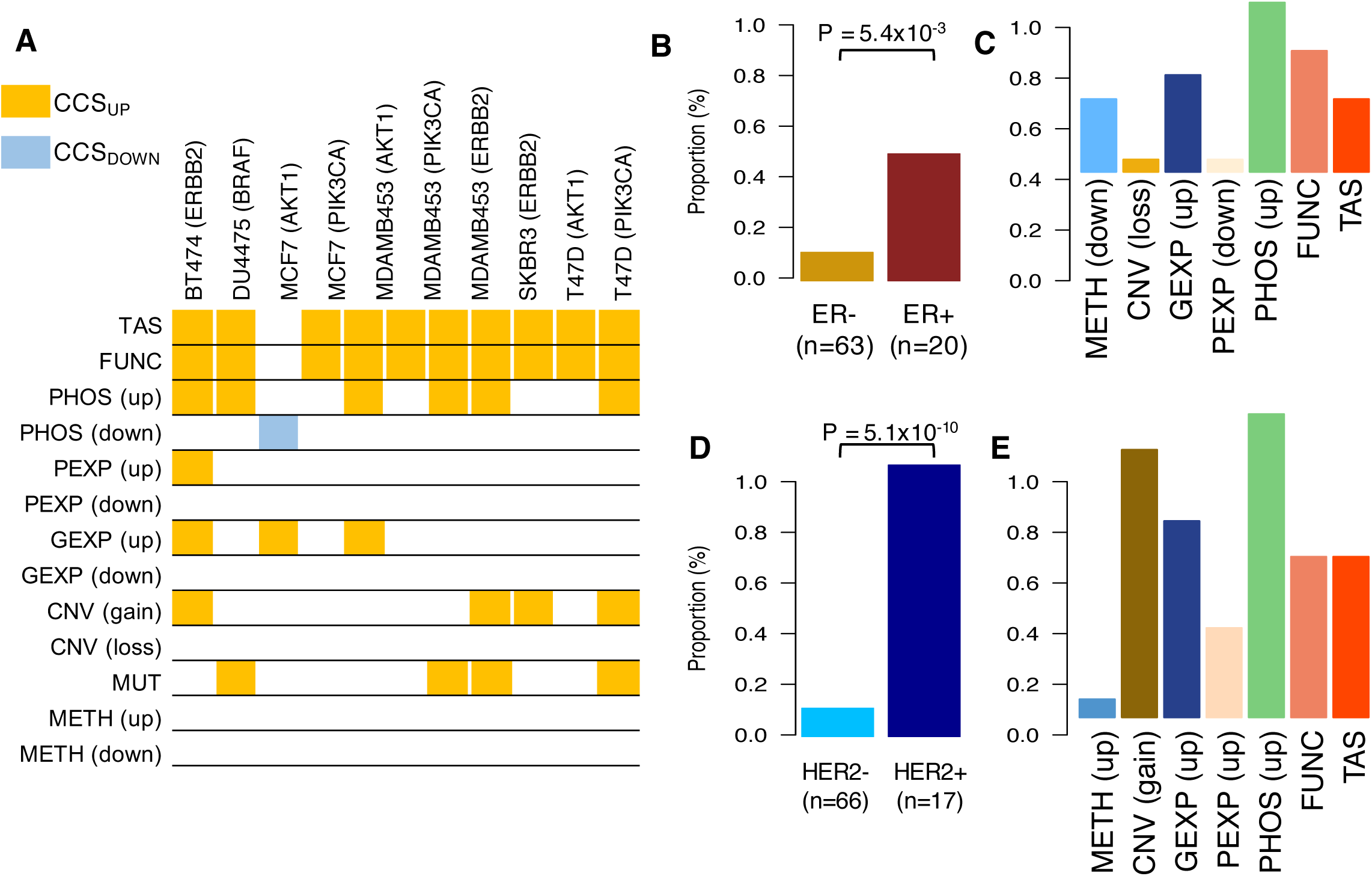
CLIP-signature of established breast cancer driver kinases (A) Subset of cell line-specific drivers that were identified as rCCS genes in this study. Highlighted entries indicate that the gene was identified as a rCCS gene in that modality. (B) Proportion of ER+ and ER- breast cancer cell lines that have ESR1 as a rCCS gene. P-value was calculated based on Fisher’s exact test. (C) The data modalities that supported the rCCS status of ESR1 and the proportion of cell lines having that evidence in the ER+ cell lines. (D) Proportion of HER2+ breast cancer cell lines that have ERBB2 as a rCCS gene. P-value was calculated based on Fisher’s exact test. (E) The data modalities that supported the rCCS status of ERBB2 and the proportion of cell lines having that evidence in the HER2+ cell lines.

We further expanded the individual cancer cell line-specific approach to identify rCCS genes within a group of cell lines. Breast cancer cell lines are conventionally categorized based on the expression levels of ER and HER2 receptors into three subtypes, indicative of their clinical characteristics(Koboldt et al., 2012; Perou et al., 2000; Van’t Veer et al., 2002) (Table S7). We therefore reasoned that the CLIP framework should be able to identify the relevant receptor proteins as rCCS genes in the cell lines belonging to these subtypes. Indeed, we observed that ER and HER2 were more frequently identified as rCCS genes in the subtype-specific cell lines (Figure 4B, D), suggesting that our data-driven approach to identifying context-specific players for each cell line is able to recapitulate the known molecular features of these cell lines. Further, upon investigating the supporting rCCS evidence from the different molecular modalities, we found that these genes had shared support at functional, gene expression and protein phosphorylation levels (Figure 4C, E). We further observed that methylation levels of ER were downregulated in a few of the rCSS-identified ER+ cell lines (Figure 4C). Similarly, in cell lines driven by ERBB2, the rCCS status was also supported by copy number gain, as it is known that ERBB2 is frequently amplified in HER2+ cell lines (Figure 4E).

As for an additional support for the data-driven discoveries enabled by CLIP, we further observed that a number of previously-reported highly expressed genes, such as GATA3 in ER+ tumors(Koboldt et al., 2012; Perou et al., 2000) and PGAP3, GRB7 and STARD3 that are frequently co-amplified with HER2(Koboldt et al., 2012; Perou et al., 2000), were also in the list of most significant rCCS genes for the ER+ and HER2+ subtypes, respectively (Table S8). Similarly, SMAD7 was identified as one of the rCCS genes in the triple negative breast cancer (TNBC) subtype (Table S8). SMAD7 is known to play a role in metastasis and epithelial-to-mesenchymal (EMT) transition, a feature is frequently exhibited by the TNBC tumors(Katsuno et al., 2018; Valcourt et al., 2005). These results suggest that the CLIP framework is able to pinpoint the established cell line and subtype-specific drivers, and also corroborate the mechanistic evidence for the genes involved in breast cancer progression. Importantly, many of these drivers would have been missed when looking at one of the studies or molecular modalities alone, rather an integrative approach was necessary to identify the robust and reproducible driver signatures. In addition to the known markers, which were used here as positive controls, the CLIP framework identified a number of novel genes for specific breast cancer subtypes (Table S8), which provide leads for future research.

### CLIP identifies ECHDC1 as a novel tumor suppressor in breast cancer

We observed that many of the known key players of breast cancer, such as BRCA1, ERBB2, ESR1, GATA3, CDH1, FOXA1, were frequently identified as rCCS genes by CLIP (Table S9), suggesting that the integrative approach is capable of ascertaining genes with relevant biological implications for breast cancer. Interestingly, we also observed frequently-selected novel rCCS genes, such as ECHDC1, SYCP2, GPX1 and MSN, whose role in breast cancer have not yet been studied extensively (Table S9). In particular, ECHDC1 was selected as the most frequent rCCS gene among 20 out of the 102 breast cancer cell lines considered in our analysis. ECHDC1 encodes an enzyme, Ethylmalonyl-CoA Decarboxylase 1, with a potential metabolite proofreading function(Linster et al., 2011). Interestingly, a previous report based on genome-wide association study implicated the genomic locus mapping to the ECHDC1 as a breast cancer risk locus in Jewish Asheknazi women(Gold et al., 2008). Notably, neither metabolic profiling nor germline genotyping data were used as part of the CLIP framework, thereby these studies provide an orthogonal support for a previously unappreciated role of ECHDC1 in breast cancer.

Our further analysis of the CLIP-Signature of ECHDC1 revealed that it is hyper-methylated in all the breast cancer cell lines in which it was identified as a rCCS gene (Figure 5A. In the same cell lines, ECHDC1 mRNA was downregulated, suggesting that ECHDC1 could be a putative tumor suppressor(Esteller, 2002). Moreover, we also observed that higher methylation levels of ECHDC1 were associated with reduced survival probability of breast cancer patients in The Cancer Genome Atlas (TCGA) dataset (P= 0.0015, log-rank test), corroborating its putative tumor suppressive role (Figure 5B). To experimentally challenge this finding, we knocked down ECHDC1 in the immortalized human MCF10A breast epithelial cell line and malignant BT-474 cell line, and embedded the cells in 3D collagen matrix (Figure 5C). The ECHDC1-depleted MCF10A cells acquired a more invasive and proliferative phenotypes, when compared to scramble control, whereas BT-474 phenotype remained unaltered after the knockdown (Figure 5C, Figure S7), further supporting the tumor suppressive role of ECHDC1 in breast cancer.

**Figure 5:**
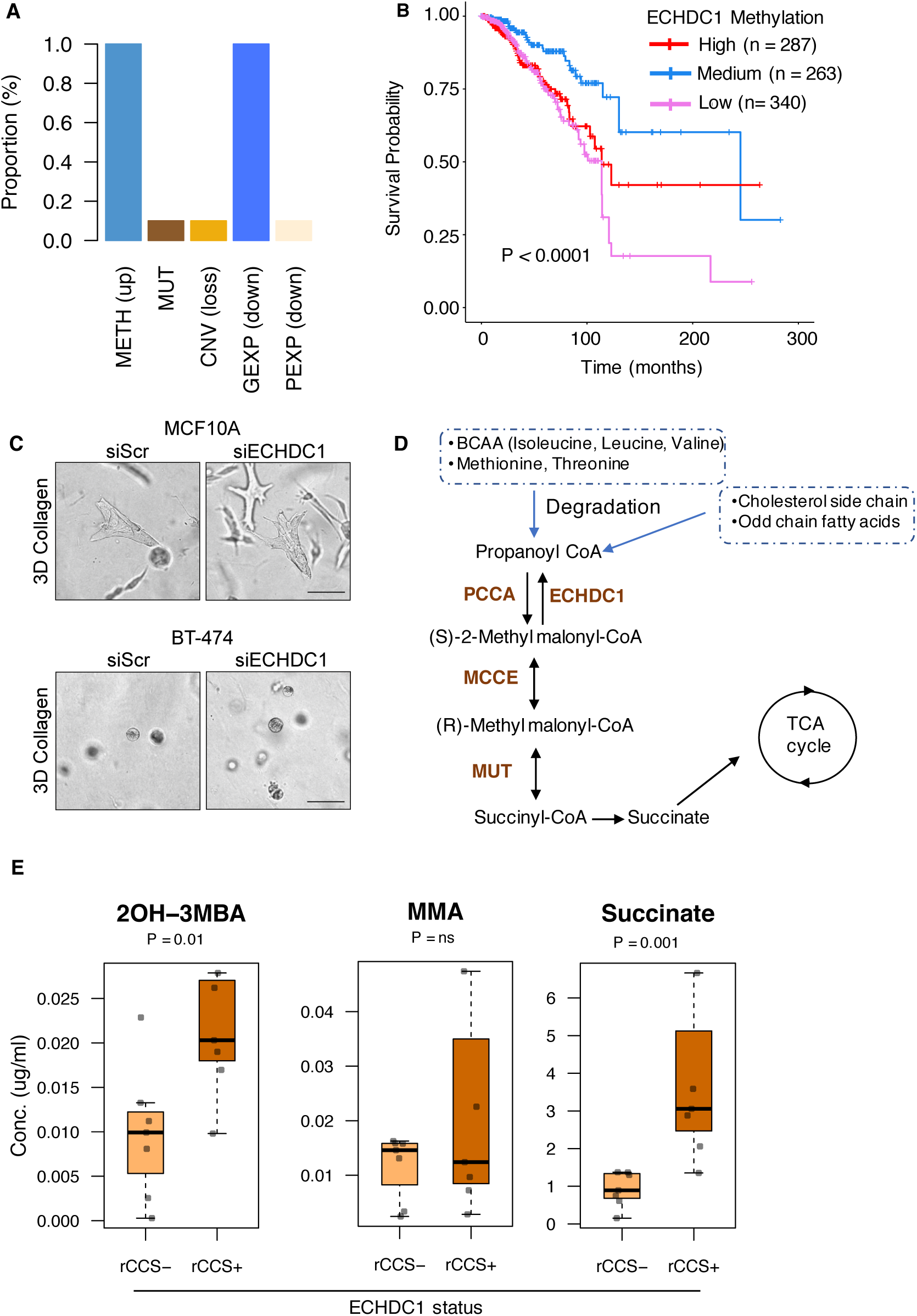
Identification of ECHDC1 as breast cancer tumor suppressor gene. (A) The CLIP-signature of ECHDC1 suggests that it was hypermethylated and down-expressed in all the breast cancer cell lines in which it was identified as rCCS gene. (B) Survival analysis based on average methylation levels of ECHDC1 in breast cancer patient tumors in the TCGA dataset. Methylation levels, quantified as beta values (*B*), were divided into 3 classes: low, medium and high, based on tertiles (< 0.34, 0.34 - 0.38, > 0.38 respectively). Patients in the high class have lower survival probability than those in the medium class (P = 0.0015; log-rank test). Overall, survival probabilities differed between the 3 classes (P = 0.00014; log-rank test). C) Representative light micrographs of ECHDC1 silenced MCF10A and BT-474 cells grown in 3D collagen for 5 d. In rCCS+ BT-474 cells, knockdown of the already downregulated ECHDC1 did not alter the cell phenotype, as expected, whereas the benign breast epithelial MCF10A cells acquired an invasively growing and more proliferative phenotype. (D) Metabolic pathway of propanoate metabolism. (E) Measured metabolite levels of intermediates in propanoate metabolism in select breast cancer cell lines with or without the ECHDC1 rCCS status (n=7 in both groups, see Suppl. Fig. S7). Statistical significance was assessed using Wilcoxon test.

To illuminate the mechanistic basis of the tumor suppressive role of ECHDC1, we investigated the metabolic pathway in which ECHDC1 is involved, namely, the propanoate metabolism (Figure 5D, Figure S8). Propanoyl-CoA is an end product of catabolism of several branched chain amino acids, and oxidation of cholesterol side chains and odd-chain fatty acids. Propanoyl-CoA is further converted to succinyl-CoA, which is oxidized and fed into the TCA cycle. We reasoned that the down-regulation of ECHDC1 in breast cancer cells could lead to alteration in the levels of intermediate metabolites resulting in tumorigenesis. Sub-sequent metabolite profiling of three such intermediate metabolites revealed that succinate and 2OH-3MBA were significantly up-regulated in the breast cancer cell lines in which ECHDC1 was identified as a rCCS gene (Figure 5E). Succinate is known to be elevated in various cancers(Dalla Pozza et al., 2020; Zhao et al., 2017), and it may potentially contribute to tumor imitation and progression through regulation of mitochondrial function, hypoxia and reactive oxygen species production.

These observations further strengthen our data-driven approach, and suggests that ECHDC1 is a novel tumor suppressor of breast cancer. This was also supported by a pathway co-regulation analysis for predicting gene function (see Materials and Methods), which suggests ECHDC1 is likely to play a role in TCA cycle and mitochondrial respiration, namely the electron transport chain, and fatty acid beta-oxidation pathway (Figure S9).

### CLIP predicts novel genetic interaction partners for cancer drivers

To further extend the applicability of our integrative meta-analysis approach, we reasoned that the CLIP framework could also identify novel and robust genetic interaction partners of cancer driver genes. Such genetic co-addictions, more specifically synthetic lethal (SL) interactions, exhibit differential dependencies in context-specific genetic backgrounds, e.g., presence or absence of a cancer driver mutation(Ashworth et al., 2011; Kaelin, 2005; Nijman and Friend, 2013). These co-addictions are often observed only in certain cell lineages, making their identification challenging in smaller-scale studies(Huang et al., 2020; Nijman and Friend, 2013).

As CLIP identifies context-specific rCCS genes in large panels of cell lines, we reasoned that any gene that is both supported by the gene dependency modality and identified frequently as a rCCS gene specifically in cancer cell lines mutated for a cancer driver, could provide a potential SL interaction partner for the driver gene. To examine this rationale for identifying context-specific and reproducible SL interaction partners, we first confirmed that CLIP was able to identify the known oncogenic addictions, such as KRAS, PIK3CA and BRAF as rCCS genes, in the specific cell lines that harbor these oncogenic driver mutations (Figure 6 A-D and Figure S10 A-B, Table S10). Cancer cell lines with such oncogenic driver mutations are known to be dependent on the same driver genes, due to oncogenic addiction(Weinstein and Joe, 2008), supporting the use of gene dependency modality in their detection. Moreover, we extended this analysis to identify also co-addiction partners of other major cancer driver genes that are also frequently mutated in specific cell contexts, and in doing so, we identified a previously reported SL interaction between ARID1A and ARID1B(Helming et al., 2014), suggesting that the approach is able to recapitulate many confirmed SL interaction partners (Table S10).

**Figure 6:**
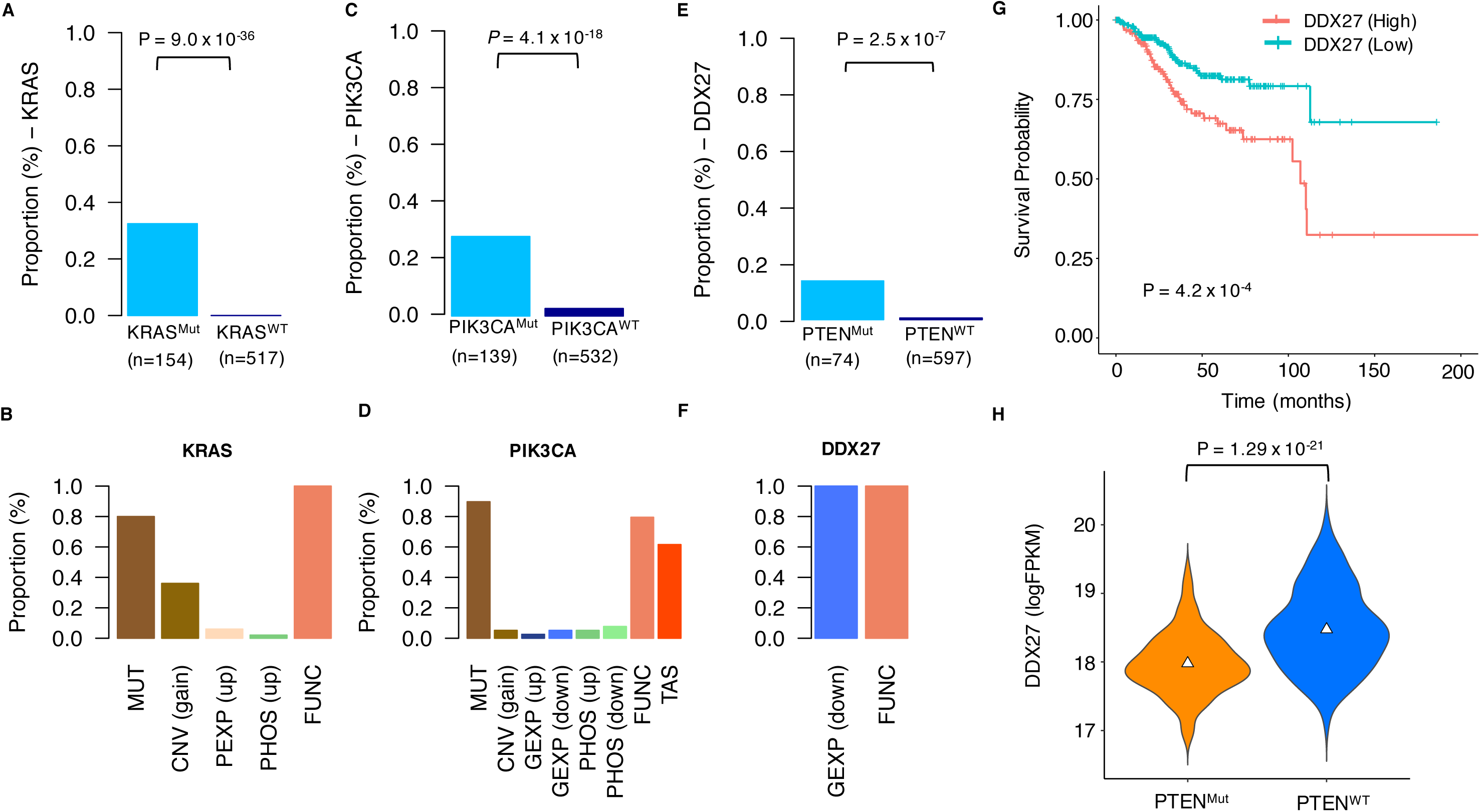
Identification of novel synthetic lethal interactions. (A) Proportion of KRAS mutated (Mut) and KRAS wild type (WT) cancer cell lines with KRAS identified as a rCCS gene. P-value was calculated based on Fisher’s exact test. (B) The modalities that support the rCCS status of KRAS and the proportion of cell lines having that evidence in the KRAS mutated cell lines. (C) Proportion of PIK3CA mutated (Mut) and PIK3CA wild type (WT) cancer cell lines with PIK3CA identified as a rCCS gene. P-value was calculated based on Fisher’s exact test. (D) The modalities that support the rCCS status of PIK3CA and the proportion of cell lines having that evidence in the PIK3CA mutated cell lines. (E) Proportion of PTEN mutated (Mut) and PTEN wild type (WT) cancer cell lines in which DDX27 was identified as a rCCS gene. P-value was calculated based on Fisher’s exact test. (F) The modalities that supported the rCCS status of DDX27 and the proportion of cell lines having that evidence in the PTEN mutated cell lines. (G) Survival analysis based on mRNA expression levels of DDX27 in patients with endometrial cancer in the TCGA dataset. Expression levels were divided into 2 classes, high (n=203 and low (n=322), based on mean expression level of DDX27 (logFPKM=18.27). Patients in the high class showed lower survival probability than those in the low class (P < 0.05; log-rank test). (H) mRNA expression levels of DDX27 in PTEN mutated (Mut, n=302) and PTEN wild type (WT, n=224) endometrial patient tumors in the TCGA dataset. P-value was calculated using Wilcoxon test.

Interestingly, we found a statistically strong evidence for DDX27 and DHX30 being SL interaction partners of the tumor suppressor PTEN (P < 0.001, Fisher’s Exact test) (Figure 6E, S10 C, Table S10). Specifically, the CLIP-Signature of DDX27 suggested that all the PTEN mutant cell lines in which DDX27 was identified as a rCCS gene had downregulated mRNA levels of DDX27, which was also essential for their growth (Figure 6F). Both DHX30 and DDX27 belong to the DEAD box nucleic acid helicase family of proteins that have been recently shown to modulate the formation of RNA molecular condensates, known as stress granules, thereby exerting a role in ribosomal translation(Fuller-Pace, 2013; Ivanov et al., 2019; Tauber et al., 2020). Recently, DDX27 was shown to have a pro-tumorigenic function in colorectal cancer(Tang et al., 2018), and we observed that it correlates with poor prognosis in endometrial cancer (Figure 6G), liver and renal cancer in the TCGA dataset (Figure S10 D). It is noteworthy that PTEN is the most frequently mutated gene in endometrial cancers (Figure S10 E), but with no drugs available for its direct reactivation. In line with the mutual exclusivity property of many SL partners,(Unni et al., 2015; Varmus et al., 2016) we also found that endometrial tumors harboring loss-of-function PTEN mutations had much lower expression levels of DDX27 (Figure 6H, P = 1.29×10^-21^, Wilcoxon test), and this property was also significant in the TCGA Pan-Cancer dataset (Figure S10F, P = 1.06×10^-19^). The mechanistic basis of this co-dependency is likely through the effects on protein synthesis and translation. The loss of PTEN induces an increased physical association of mTORC2 and ribosome, which drives cancer growth while making the cells stress-prone and vulnerable to apoptosis(Keniry and Parsons, 2008; Zinzalla et al., 2011). Downregulation of transcripts of RNA helicases together with a compounded loss of its activity could severely limit the ability of cells to cope with the induced stress by preventing the formation of RNA:RNA aggregates, thereby making the cells apoptosis prone(Ivanov et al., 2019; Tauber et al., 2020).

## Discussion

In this study, we first provided a comprehensive and quantitative view of the reproducibility of multiple data modalities of cancer cell lines by means of a systematic and non-parametric correlation analyses of molecular profiles of identical cell lines profiled in different laboratories. In particular, we found out that the profiles based on genomic technologies, such as transcriptome and copy number alterations (CNA), are highly consistent across the labs (Figure 1D). This is rather expected, given the robustness and maturity of the sequencing and hybridization techniques. Contrary to our expectations, we found that that the consistency of point mutational profiles was relatively low, even though the correlation difference in point mutation profiles between identical cell lines compared to non-identical cell lines was still highly significant (Figure 1D). This could be partly attributed to the correlation metric used for evaluating reproducibility (Matthew’s or Spearman’s), as we observed also a decrease in correlation of binarized CNV calls compared to continuous copy number intensity calls (Figure S11).

While previous studies(Barretina et al., 2012; Ben-David et al., 2018) have also compared the consistency of mutational profiles between different laboratories, their approaches are different from ours. Barretina et al.(Barretina et al., 2012) compared the agreement between cancer cell lines based on mutational frequency of genes in CCLE with patient tumor based mutational frequency from the COSMIC project. Likewise, Ben-David et al.(Ben-David et al., 2018) compared the allelic fraction of somatic variants in 106 cell lines common between CCLE and GDSC datasets. Interestingly, they observed that 10-90% of non-silent mutations identified in one dataset were not identified in the other, suggesting variability in the point mutational profiles of identical cell lines. It is also noteworthy that we observed a wide variation in the correlation of CNA profiles across the laboratories, suggesting that cell lines might have undergone clonal selection at different research sites. Such clonal divergence between identical cell lines has profound implications for the conclusions drawn from experimental assays performed on cultured cancer cell lines(Ben-David et al., 2018). We also investigated the various factors that contributed to the variation in the reproducibility estimates and found, for instance, significant discrepancies in the proteomic profiles between TMT-labeled and non-labeled MS techniques.

In the next phase, we developed a non-parametric integrative meta-analysis framework to identify robust molecular determinants, unique to an individual cancer cell line, that are shared among multiple data modalities and studies. One of the limitations for the integrative analysis was that the different research sites have profiled different panels of cell lines, making it challenging to derive robust estimates for every cell line. Nevertheless, we found extensive profiling information of breast cancer cell lines across multiple sites, which served here a purpose for evaluating the performance of our approach. We demonstrated that our reproducibility-based integrative framework was able to identify well-established breast cancer driver genes as robust cancer cell-line specific (rCCS) genes using the available omics data from the set of breast cancer cell lines. Further, we extended this bottom-up approach for the identification of individual CCS genes, and demonstrated that this approach also recapitulates the known drivers at a broader sub-group level; such as the well-established ER+ and HER2+ sub-types of breast cancer.

As an application case, our non-parametric CLIP framework identified a novel driver gene, ECHDC1, with a hitherto unknown tumor suppressive role in breast cancer. We demonstrated the tumor suppressive role of ECHDC1 and highlighted a possible mechanism by which ECHDC1 may contribute to breast tumor growth. Specifically, we showed that the level of succinate increases in breast cancer cells with hyper-methylated and lowly-expressed ECHDC1. Moreover, it has been shown previously(Dalla Pozza et al., 2020; Zhao et al., 2017) that higher levels of succinate are associated with tumorigenesis and cancer progression via dysregulation of mitochondrial function, a hypoxic environment and production of ROS, which all have established roles in the etiology of cancer development. Additionally, several tumors are also known to have inactivating mutations in the SDH(Dalla Pozza et al., 2020), succinate dehydrogenase, the enzyme that processes succinate and feds in to the TCA cycle. We further applied the CLIP framework to identify context-specific synthetic lethal relationships with well know cancer driver genes in a wider panel of cancer cell lines from multiple cell types. In addition to capturing known aspects of oncogenic dependency, we captured a novel co-dependency relationship between PTEN loss-of-function mutations and RNA helicase enzymes DDX27 and DHX30. The cell lines that had a higher prevalence of loss-of-function mutations in PTEN had lower expression of DDX27, and this mutually exclusive genetic interaction was particularly strong in endometrial cancers.

In summary; firstly, this study provides to date the most comprehensive perspective on the reproducibility of molecular profiles of cancer cell lines, and delineates specific factors that contribute to the consistency that should be considered in future studies. Secondly, to provide solution to the sub-optimal consistency, we developed an integrative meta-analytic framework for leveraging robust and reproducible signal from various modalities of molecular profiles that also accounts for the observed variation between datasets generated at different laboratories. Finally, this study also demonstrates the potential of such integrative approaches for identification of novel molecular features having a confirmed role in breast cancer. We expect this approach will lead to many more exciting discoveries once more profiling data becomes available also from other cancer types.

## Acknowledgements

We thank the authors of the various projects considered in the study for making their data publicly available. We also thank Dr. Anil Giri for his help with the methylation data analysis, and Dr. Vidya Velagapudi for her help with designing the metabolite assays.

## Availability of data and materials

Processed datasets used in the study are publicly available at https://github.com/jaiswal-alok/clip-meta

## Authors’ contributions

AJ conceived and conducted the study. AJ analyzed the data and drafted the manuscript. PG, EP, ST and NN contributed to cell line laboratory experiments. NS contributed to metabolite measurements. ZT contributed to data analysis. KL contributed to overseeing the cell line experiments. KW critically reviewed the manuscript. TA supervised the study and revised the manuscript. All authors read and approved the final manuscript.

## Consent for publication

Not applicable.

## Ethics approval and consent to participate

Not applicable.

**Figure S1:**
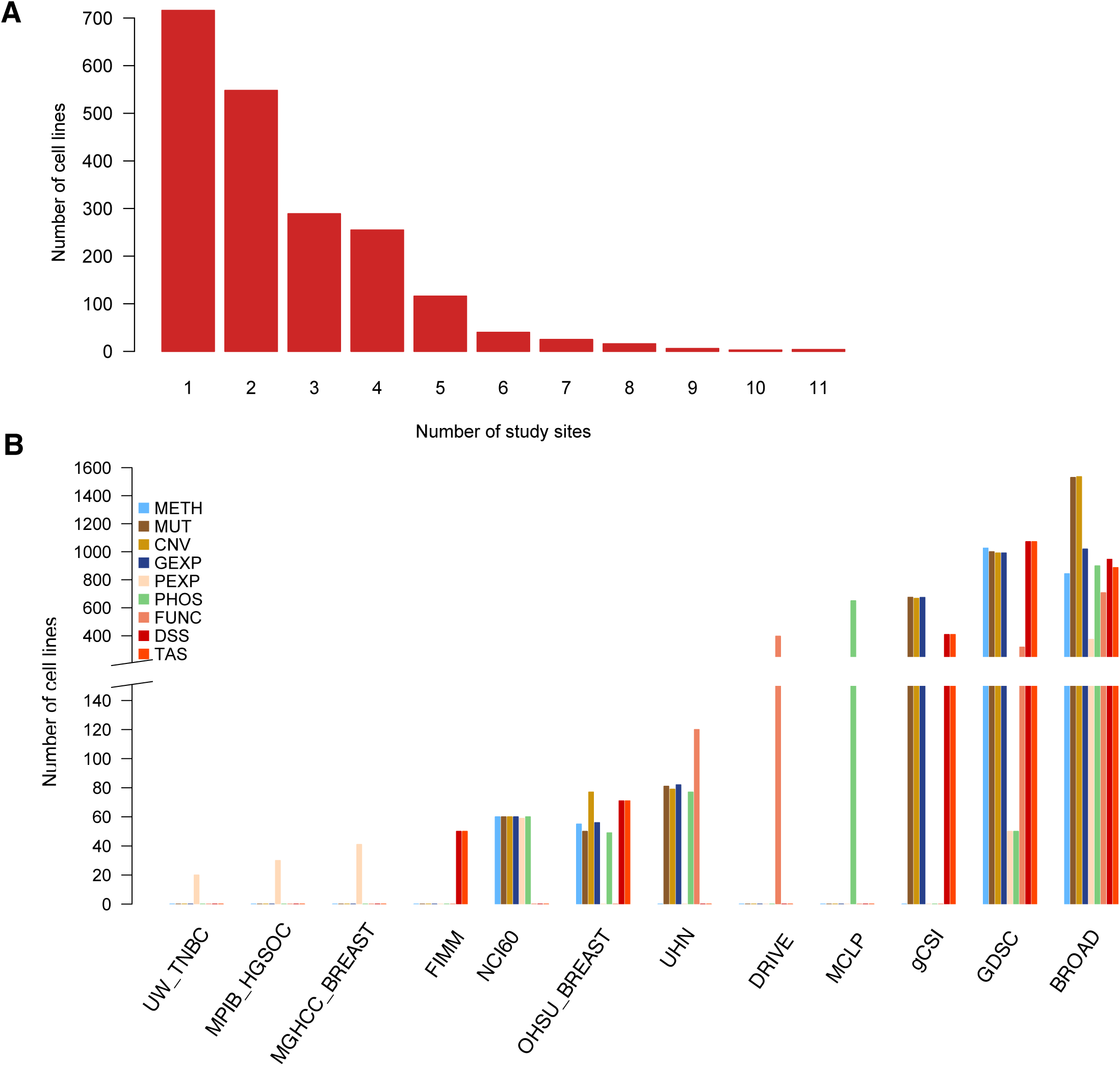
(A) The number of cell lines having available data from the 12 different research sites that were considered in the study. (B) The number of cell lines having available data at each research site by data modality type. METH = Gene promoter methylation, MUT = Gene point mutation, CNV = Copy number variation, GEXP = Gene mRNA expression, PEXP = Protein expression, PHOS = Protein phosphorylation, FUNC = Gene dependency, DSS = Drug sensitivity, TAS = Protein addiction.

**Figure S2:**
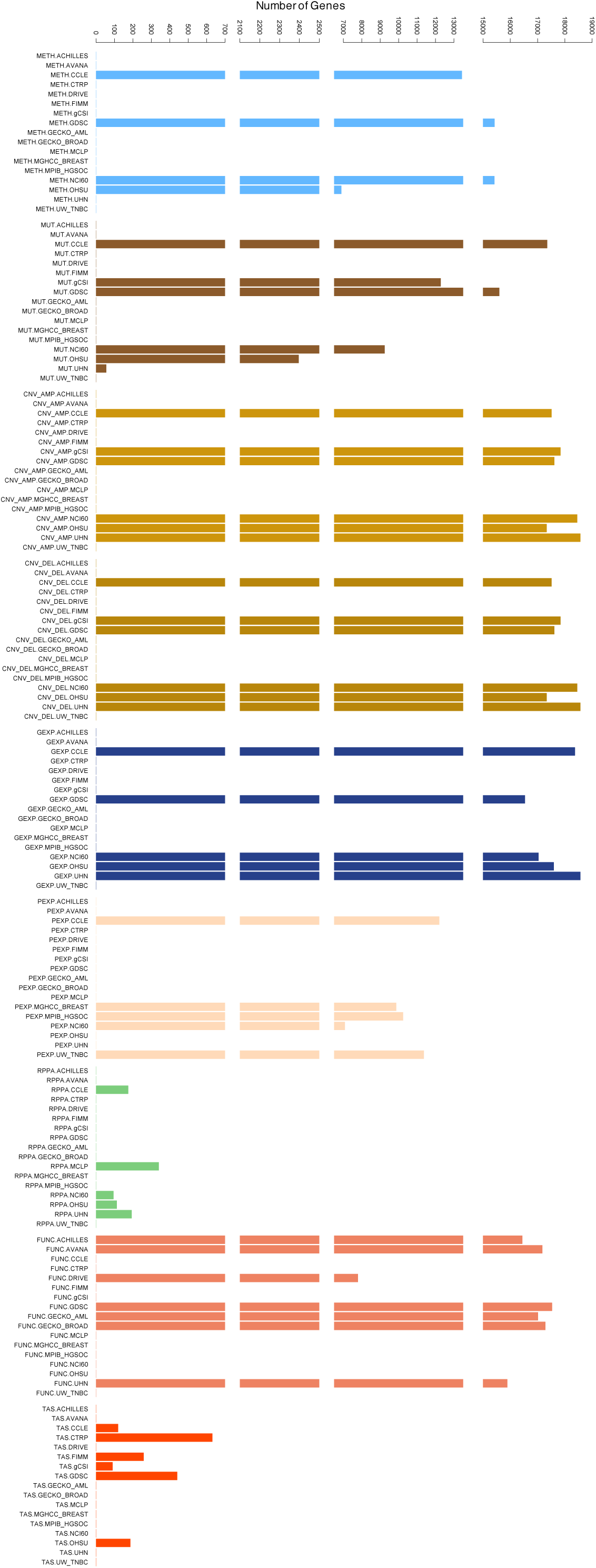
Breakdown of number of genes for which data was available for each modality at each research site. For CNV modality, the datasets were categorized into amplifications and deletions, namely CNV_AMP and CNV_DELs which were included in the CLIP framework, METH = Gene promoter methylation, MUT = Gene point mutation, CNV = Copy number variation, GEXP = Gene mRNA expression, PEXP = Protein expression, PHOS = Protein phosphorylation, FUNC = Gene dependency, DSS = Drug sensitivity, TAS = Protein addiction.

**Figure S3:**
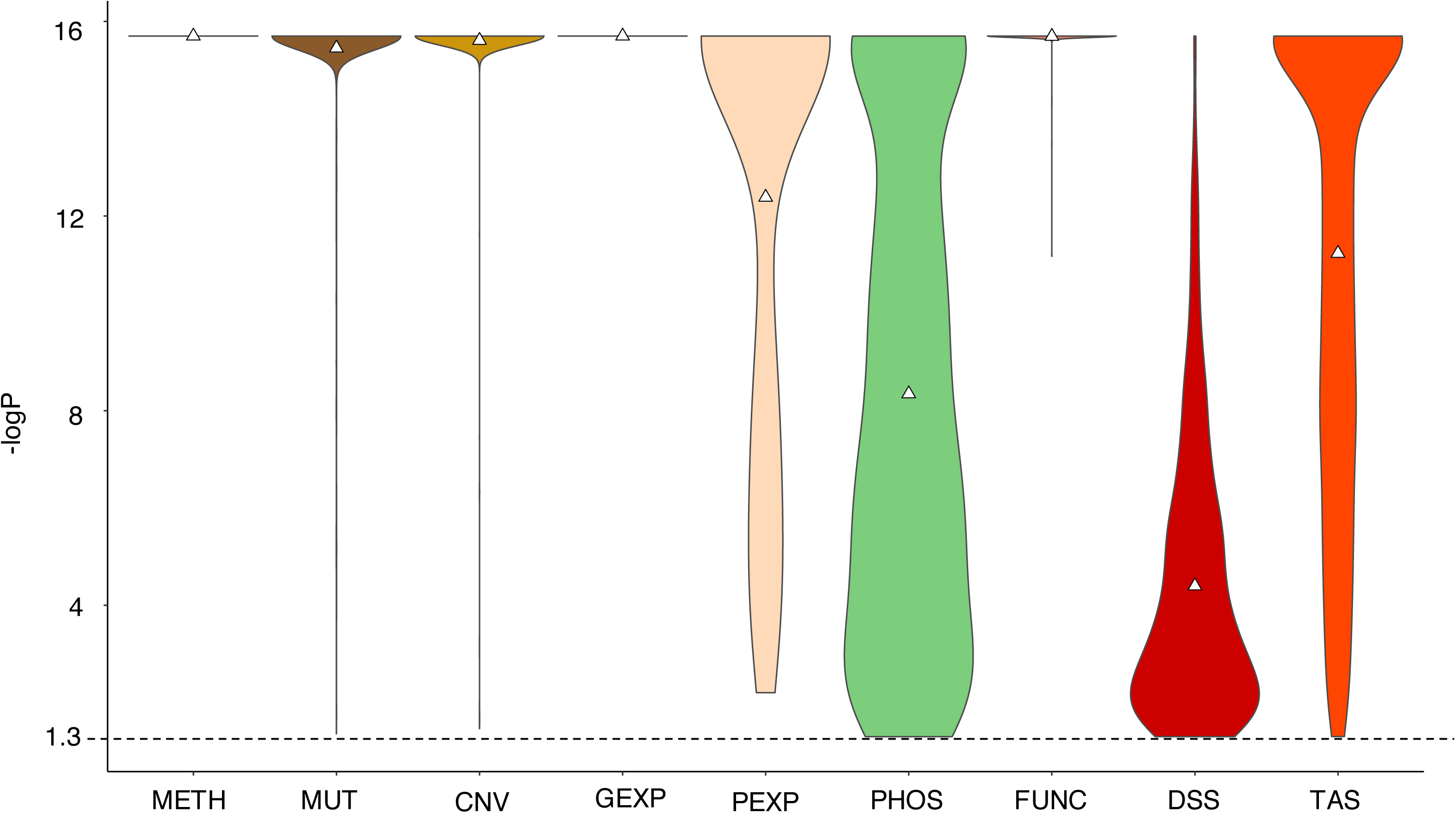
Negative logarithm of the P-values (-logP) calculated for all the correlations among identical cell lines between research sites to study the effect of sample size (N_g_) on the estimated correlations. P-values < 2.0 x 10^-16^ were set to 2 x 10^-16^. The dotted line indicates the significance cut-off of P=0.05. The triangle symbol indicates mean.

**Figure S4:**
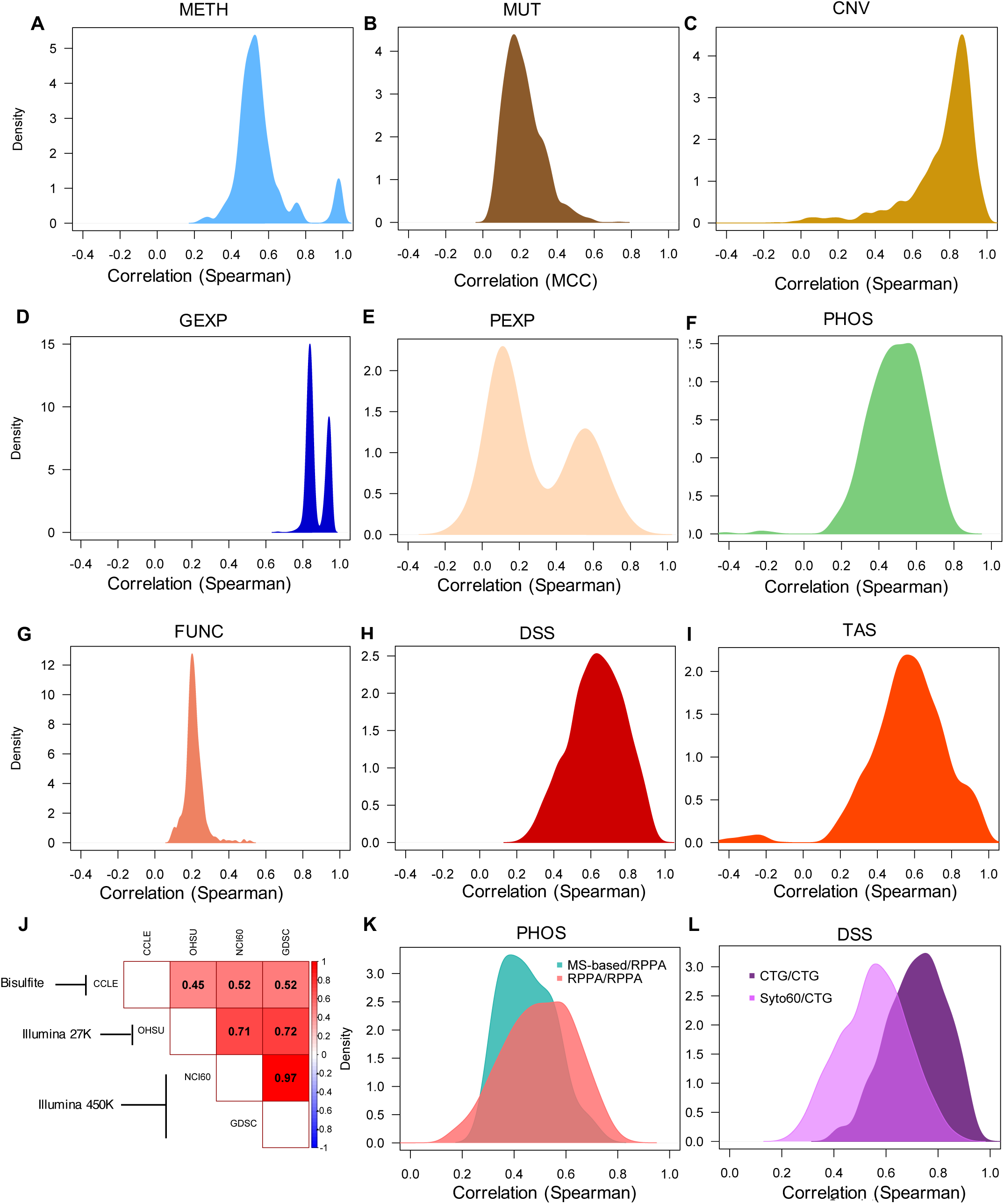
(A-I) Density plot of Spearman correlation coefficients of identical cell lines between multiple study sites for each data modality. The presence of bi-modality indicates a major technical or biological factor contributing to the mixed distributions. (J) Average spearman correlation of methylation profiles of cancer cell lines generated at different study sites (columns) and with different technical platforms (rows). (K) Density plot of Spearman correlation estimates between average gene phosphorylation profiles of cancer cell lines generated at different study sites using RPPA and MS-based technologies. (L) Density plot of the Spearman correlation estimates between drug sensitivity profiles of cancer cell lines generated at different study sites using CellTiterGlo (CTG) and Syto60 assays.

**Figure S5:**
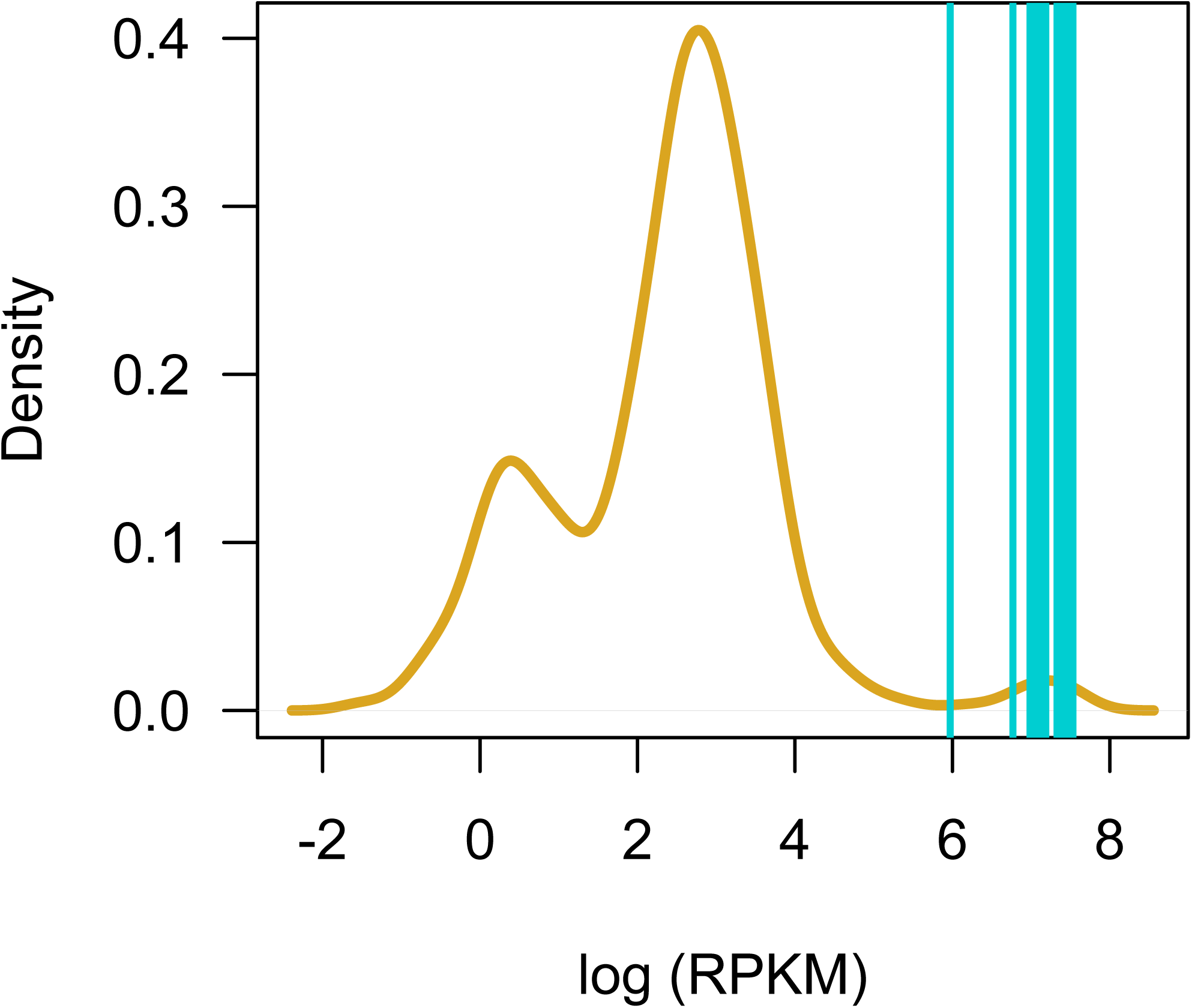
mRNA expression levels of ERBB2 gene in all cancer cell lines profiled in the CCLE dataset is plotted as a density distribution. Expression levels of ERBB2 in breast cancer cell lines that are known to be HER2+ are marked by the turquoise colored vertical lines. As the expression levels of ERBB2 in HER2+ breast cancer cell lines are towards the right tail, it can be considered that ERBB2 is a CCS gene in these cell lines.

**Figure S6:**
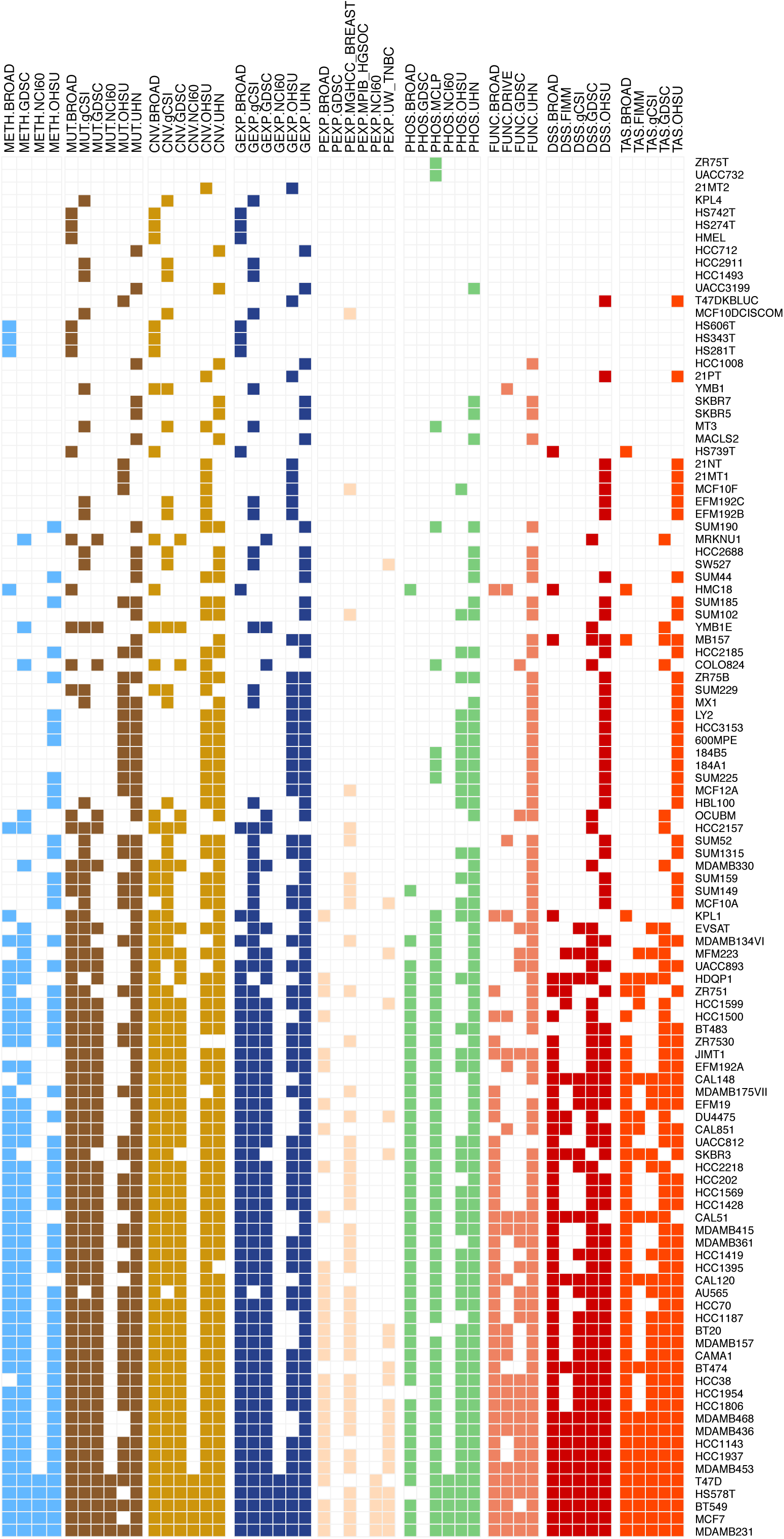
Data availability of breast cancer cell lines (columns) for each of the modality type from multiple research sites (rows).

**Figure S7:**
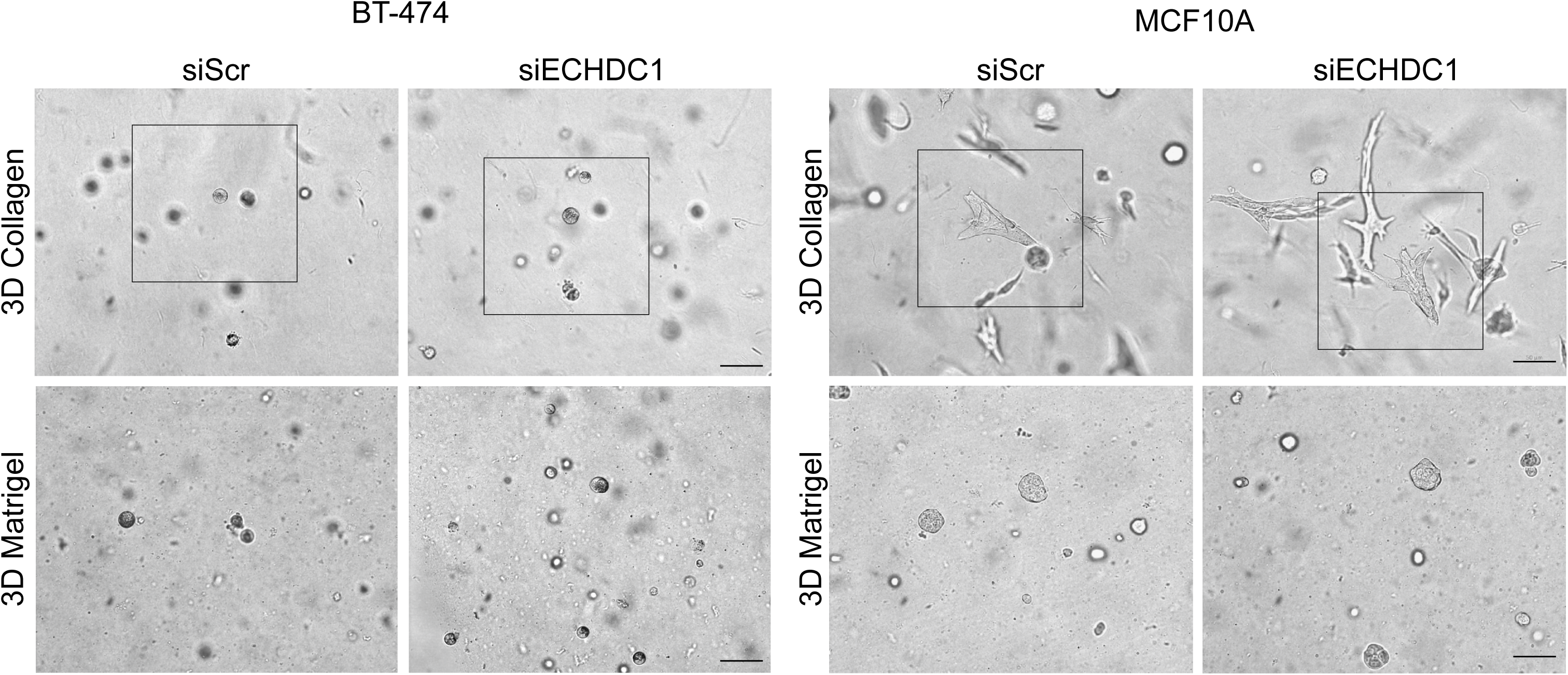
Representative light micrographs of 3D collagen and Matrigel cultured BT-474 and MCF10A cells upon ECHDC1 transient knockdown. Cells were grown in 3D for 5 days after the knockdown. Marked areas represented in Figure 5. Scale bar: 50 µM.

**Figure S8:**
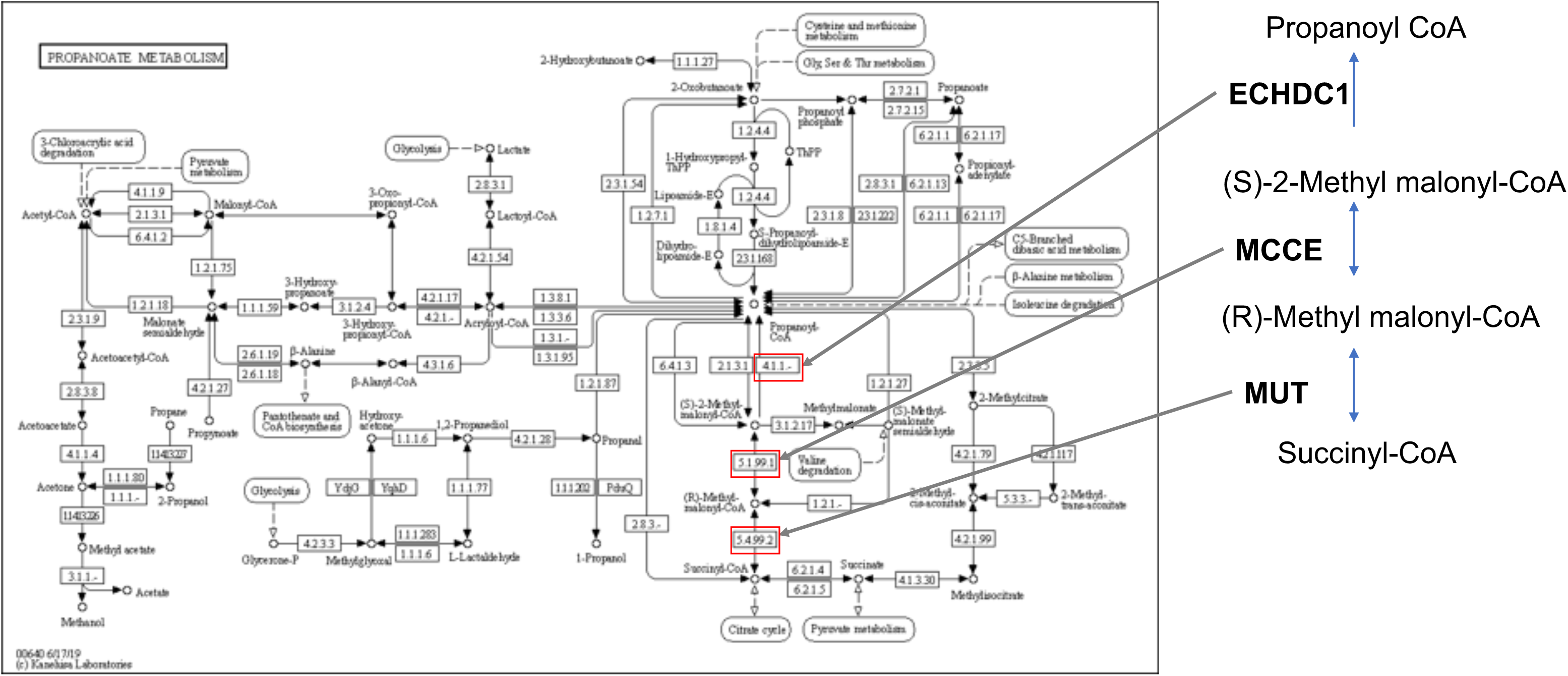
Propanoate metabolism based on the KEGG database. ECHDC1 and other genes in the pathway are highlighted.

**Figure S9:**
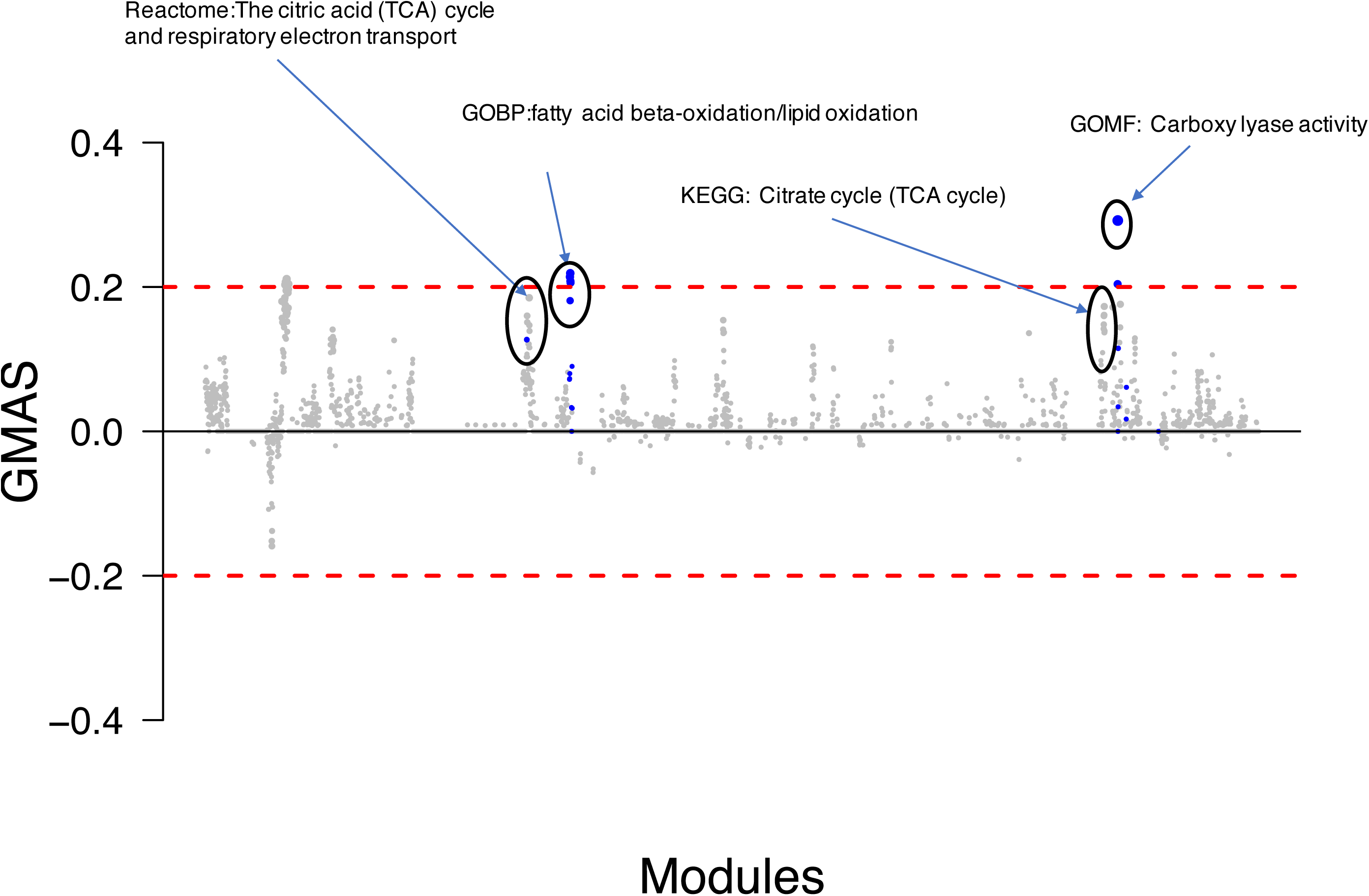
Gene Module Association Score (GMAS) for modules enriched for co-expression with ECHDC1, calculated based on Gene-Module Association Determination (G-MAD) algorithm, implemented in the GeneBridge toolkit. For details, see Table S3.

**Figure S10:**
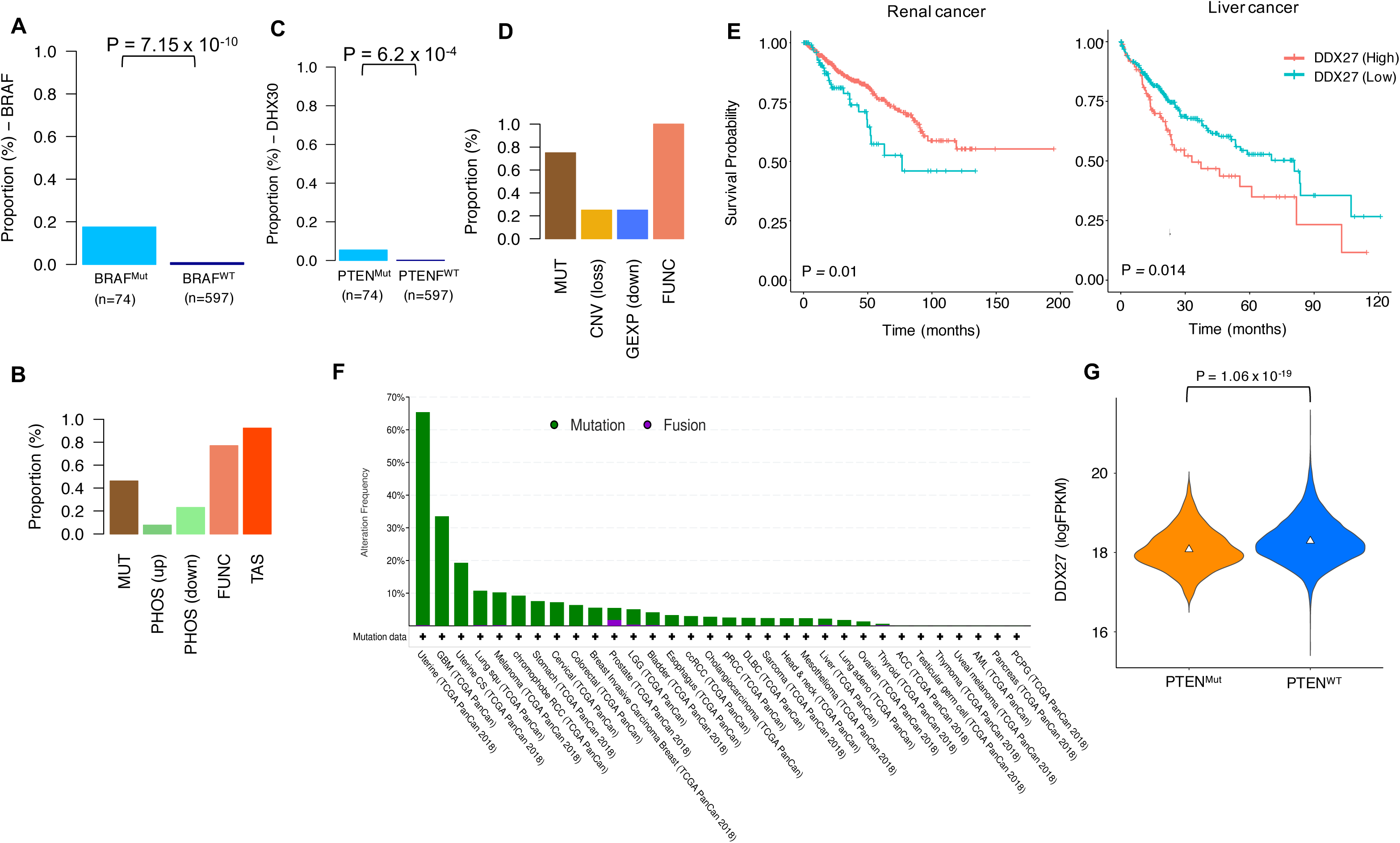
(A) Proportion of BRAF mutated (Mut) and BRAF wild type (WT) cancer cell lines with BRAF identified as a rCCS gene. P-value was calculated based on Fisher’s exact test. (B) The modalities that supported the rCCS status of BRAF and the proportion of cell lines having that evidence in the BRAF mutated cell lines. (C) Proportion of PTEN mutated (Mut) and PTEN wild type (WT) cancer cell lines in which DHX30 was identified as a rCCS gene. P-value was calculated based on Fisher’s exact test. (D) The modalities that supported the rCCS status of DDX27 and the proportion of cell lines having that evidence in the PTEN mutated cell lines. (E) Survival analysis of mRNA expression levels of DDX27 in patients with endometrial cancer, liver cancer and renal cancer in the TCGA dataset. Expression levels were divided into 2 classes, high and low, based on mean expression levels of DDX27 (logFPKM=18.27). Patients in the high class showed lower survival probability than those in the low class (P < 0.05; log-rank test). (F) Breakdown of mutational frequency of PTEN by cancer tissue type in TCGA Pan-cancer dataset, performed in cBioPortal (G) mRNA expression levels of DDX27 in PTEN mutated (Mut, n=582) and PTEN wild type (WT, n=7012) in TCGA dataset of all epithelial cancer types. P-value was calculated using Wilcoxon test.

**Figure S11:**
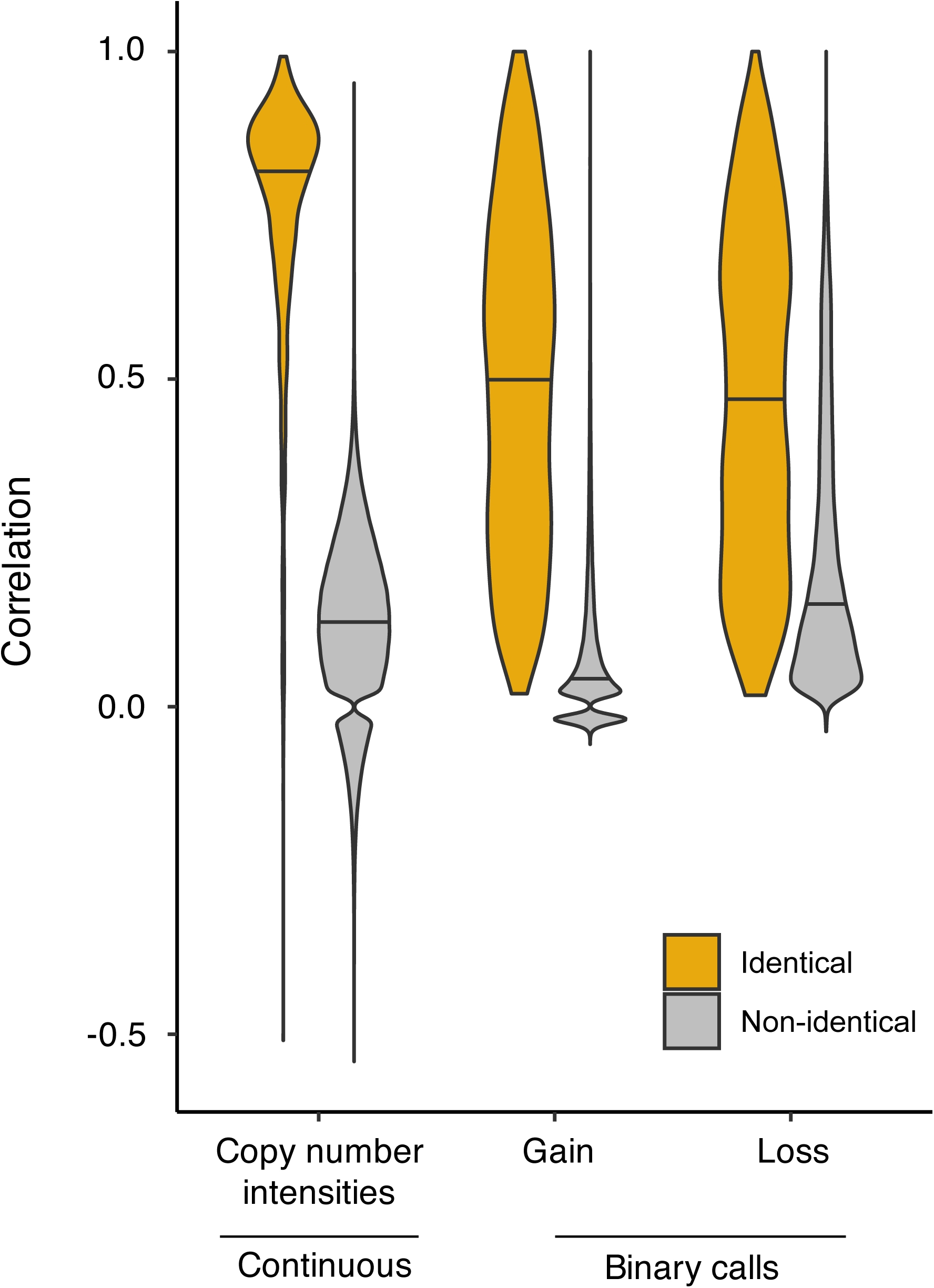
Correlation of the different types of copy number profiles. Spearman correlation was calculated between identical and non-identical cell lines between datasets from various research sites for continuous copy number calls. For binarized CNV calls of gains and losses, correlation was estimated by Matthew’s correlation coefficient.

